# Nanobodies recognizing conserved hidden clefts of all SARS-CoV-2 spike variants

**DOI:** 10.1101/2021.10.25.465714

**Authors:** Ryota Maeda, Junso Fujita, Yoshinobu Konishi, Yasuhiro Kazuma, Hiroyuki Yamazaki, Itsuki Anzai, Keishi Yamaguchi, Kazuki Kasai, Kayoko Nagata, Yutaro Yamaoka, Kei Miyakawa, Akihide Ryo, Kotaro Shirakawa, Fumiaki Makino, Yoshiharu Matsuura, Tsuyoshi Inoue, Akihiro Imura, Keiichi Namba, Akifumi Takaori-Kondo

**Author notes:** These authors contributed equally (Ryota Maeda, Junso Fujita). Correspondences (A.I.); (Ke.N.); (A.T.-K.).

## Abstract

We are in the midst of the historic coronavirus infectious disease 2019 (COVID-19) pandemic caused by severe respiratory syndrome coronavirus 2 (SARS-CoV-2). Although countless efforts to control the pandemic have been attempted—most successfully, vaccination^1–3^—imbalances in accessibility to vaccines, medicines, and diagnostics among countries, regions, and populations have been problematic. Camelid variable regions of heavy chain-only antibodies (VHHs or nanobodies)^4^ have unique modalities: they are smaller, more stable, easier to customize, and, importantly, less expensive to produce than conventional antibodies^5, 6^. We present the sequences of nine alpaca nanobodies that detect the spike proteins of four SARS-CoV-2 variants of concern (VOCs)—namely, the alpha, beta, gamma, and delta variants. We show that they can quantify or detect spike variants via ELISA and lateral flow, kinetic, flow cytometric, microscopy, and Western blotting assays^7^. The panel of nanobodies broadly neutralized viral infection by pseudotyped SARS-CoV-2 VOCs. Structural analyses showed that a P86 clone targeted epitopes that were conserved yet unclassified on the receptor-binding domain (RBD) and located inside the N-terminal domain (NTD). Human antibodies have hardly accessed both regions; consequently, the clone buries hidden crevasses of SARS-CoV-2 spike proteins undetected by conventional antibodies and maintains activity against spike proteins carrying escape mutations.

---

We have been in the historic pandemic for about two years (2019-2021): numbers of confirmed deaths and infections still increase on a weekly basis. Researchers have developed many antibodies and nanobodies as weapons to fight against COVID-19^8–11^; however, emerging SARS-CoV-2 variants of concern (VOCs) that carry mutations in the spike often escape the immune system and therapeutic antibodies^12–14^. These antibodies also serve as armuor and shields, allowing us to survey infected individuals and monitor circumstances via antigen test kits, enzyme-linked immunosorbent assays (ELISAs), dot blotting, and so forth. However, accessibility to diagnostic and monitoring kits is imbalanced worldwide because antibody-producing cells (e.g., hybridoma cells and engineered mammalian cells) are, in many cases, dominated by licenced companies that incur high costs of research, development, production, and distribution. Thus, we tried to create easily accessible and broadly neutralizing antibodies for SARS-CoV-2 VOCs.

Although many antibodies and nanobodies against the spike, especially the receptor-binding domain (RBD), have been established, those that are broadly neutralizing and detect all current variants have yet to be development. Few anti-SARS-CoV-2 spike *nanobodies* have been applied for immune assays, such as ELISAs and lateral flow and especially membrane dot blotting assays^15^. Here, we provide a panel of broadly neutralizing nanobodies that detect four SARS-CoV-2 VOCs. Our structural analyses show that two clones—P17 and P86—capped the receptor-binding domain (RBD) regardless of their up or down conformations. The most potent clone, P86, bridged a gap (Fig. 1a,b) between the down conformation of the RBD and the N-terminal domain (NTD) of a neighbouring protomer. The epitopes on the RBD recognized by the two clones partially overlapped; in particular, the epitope recognized by P17 shifted away from the L452R mutation observed in the SARS-CoV-2 delta variant spike.

**Fig. 1.**
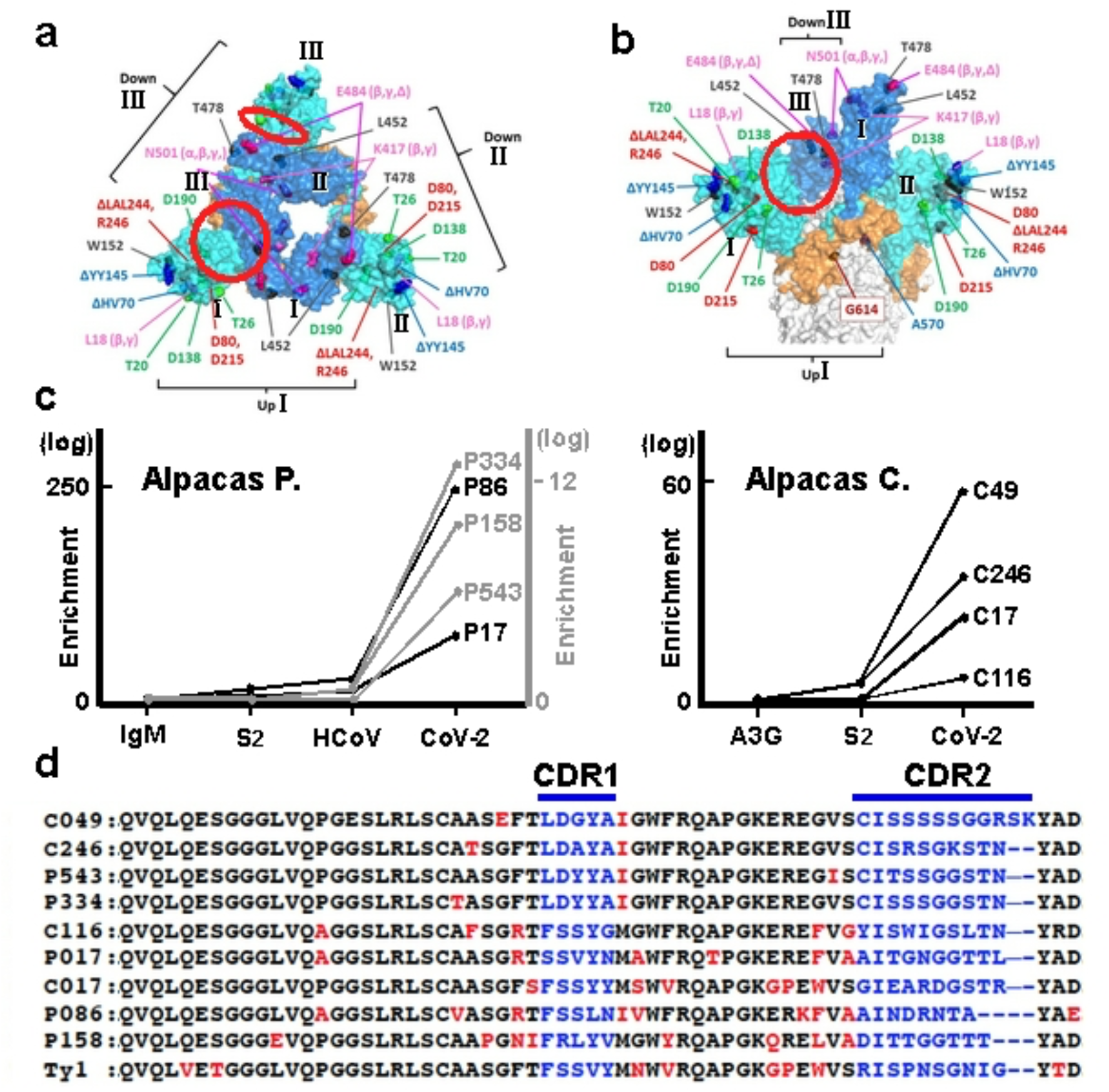
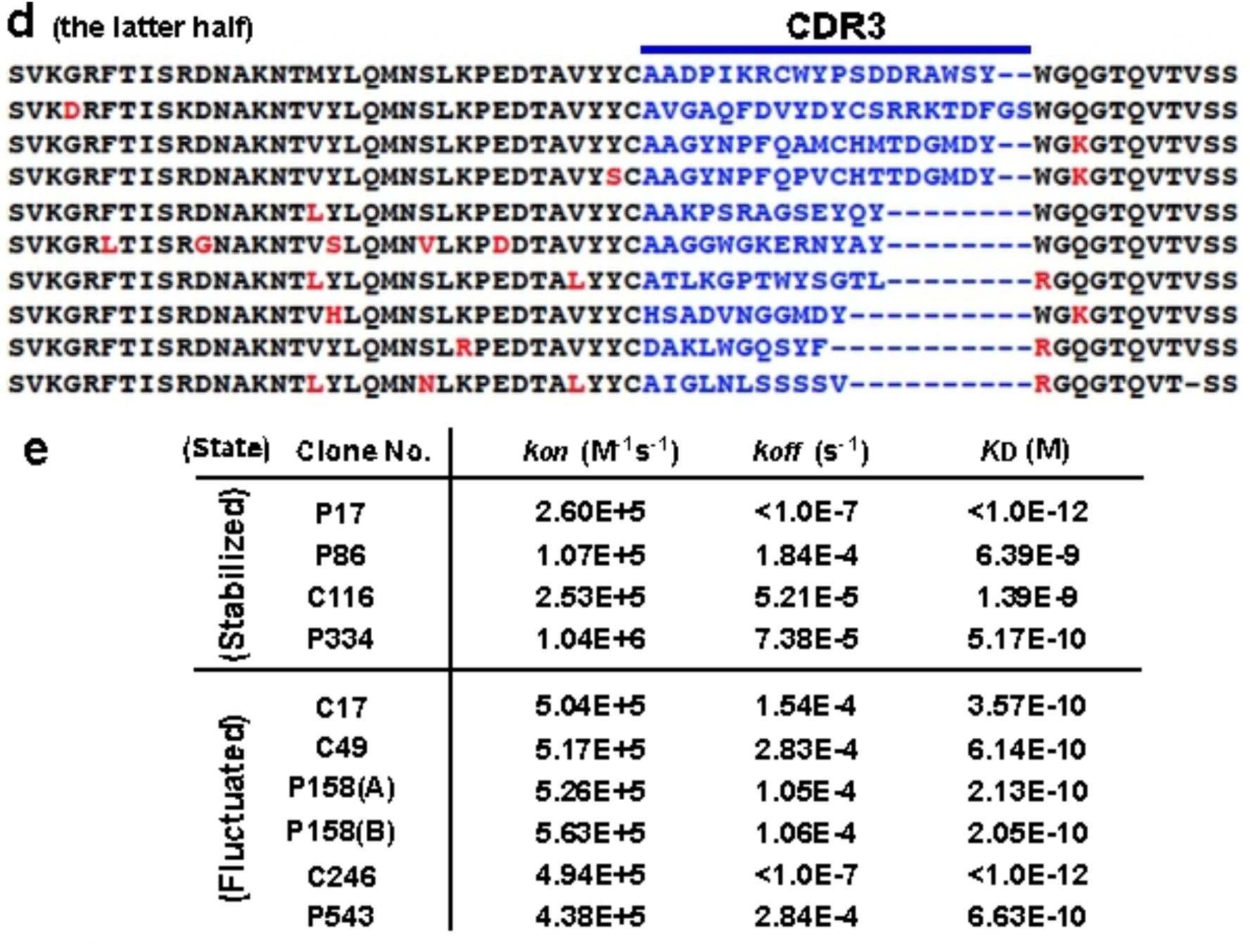
Structural features of the SARS-CoV-2 spike and the panel of nanobodies. **a,b**, Side (**a**) and top (**b**) views of the 1-up+2-down conformation of the SARS-CoV-2 spike trimer^50^ (PDB entry: 7KRR); domains of each protomer numbered I (up), II (down), and III (down). The NTD, the RBD, and the C-terminal domain (CTD) of the SARS-CoV-2 S1 domain are coloured cyan, blue, and orange, respectively; mutations and deletions studied here are indicated; clefts between the down-RBD and the NTD of the neighbouring protomer are highlighted with red circles. **c**, Graphs showing existence ratios (two different scales drawn black and grey)—*P* values in log_10_ are plotted (see Methods)—after panning with IgM; APOBEC3G (A3G)^26^; the S2 domain (S2); the HCoV-OC43 (HCoV) spike^27^; and the SARS-CoV-2 spike (CoV-2). **d**, Sequences of nine clones and that of Ty1 are compared^36^: changed amino acids in the framework regions are red; the variable complementary-determining regions (CDRs) are shown in blue. **e**, Calculated binding kinetics between each nanobody and the SARS-CoV-2 spike from sensorgrams (Extended Data Fig. 2c,d) are summarized. The upper four clones were assayed in a high-salt buffer (“stabilized” spike); the others were assayed in an anionic buffer (“fluctuated” spike).

## Purifying a SARS-CoV-2 spike protein retaining the furin cleavage site

The SARS-CoV-2 spike is one of the main targets for human antibodies^16, 17^. Its trimer complex is expressed on the surface of the virion and plays a role in fusion with host cells expressing the binding partner angiotensin-converting enzyme II (ACE2)^18, 19^. The most featured alteration of the SARS-CoV-2 spike compared to other coronavirus spikes is the generation of a furin cleavage site (-R-R-A-R^685^-: furin protease hydrolyses the C-terminal peptide bond of R^685^)^20^ located directly between the S1 (residues 1-685) and S2 (residues 686-1,213) domains^17, 21^. Because it was difficult to obtain the extracellular domain of SARS-CoV-2 spike with the furin site from the supernatant of human embryonic kidney (HEK) cells, we attempted to purify it from cell lysate. We obtained a large complex of SARS-CoV-2 spike with detergents used in crystallizing membrane proteins (Extended Data Fig. 1a-c, and see Methods)^22^. As reported previously, although peaks were broad, buffers containing high salt and detergent concentrations maintained the SARS-CoV-2 spike trimer assembly (Extended Data Fig. 1b)^23^. We noticed that a high salt concentration was necessary to stabilize the SARS-CoV-2 spike complexes (Extended Data Fig. 1b)^24^.

## Immunization of alpacas and selection of nanobodies

To obtain spike-specific nanobodies, we immunized two alpacas with the purified whole extracellular domains of the SARS-CoV-2 spike complex from cell lysates^25^ (Extended Data Fig. 1c, and see Methods). Serum titres of alpacas for SARS-CoV-2 spike increased and reached a plateau, indicating that both alpacas were effectively vaccinated (Extended Data Fig. 1d). We performed one round of biopanning with different proteins: the whole extracellular domain, the S2 domain alone of the SARS-CoV-2 spike, or the whole extracellular domain of the seasonal cold coronavirus HCoV-OC43 spike^26, 27^ (see Methods). After deep-sequencing nanobody-coding genes of each panned sublibrary, we identified nine clones that were significantly enriched in sublibraries panned with the whole extracellular domain of SARS-CoV-2 compared with those panned with the others^28^ (Fig. 1c,d, and see Methods). We expressed these clones in bacteria and purified them as C-terminally His-tagged dimers (Extended Data Fig. 2a,b); we then characterized their avidity, applicability, and neutralizing activity, as described below.

## Kinetics and detectability of nanobody binding

First, we measured the binding kinetics of these clones for the whole extracellular domain of SARS-CoV-2 spike, which was the same construct that was used for immunization, via biolayer interferometry (Fig. 1e). We noticed that the salt concentrations in the kinetic assay buffers affected the avidities of the clones. Of note, high salt concentrations were necessary to stabilize the spike complex (Extended Data Fig. 1b). Four clones (P17, P86, C116, and P334) showed high avidity for the SARS-CoV-2 spike complex in hypertonic buffer; the others (C17, C49, P158, C246, and P543) showed high avidity in hypotonic buffer (Extended Data Fig. 2c,d).

Second, we checked these nanobodies via microscopy observation using HEK cells expressing the SARS-CoV-2 spike. Seven clones stained near the cell surface; two clones (C49 and C246) stained only an intracellular region (Extended Data Fig. 3a,b). Third, flow cytometric analysis supported this observation: the seven clones detected the spike protein on the cell surface, but the C49 and C246 clones did not (Extended Data Fig. 3c). These results suggested that each clone recognizes the SARS-CoV-2 spike protein in a status-dependent manner.

## Sensitivities of the clones for the RBD mutations of VOCs

The World Health Organization (WHO) has declared four strains to be VOCs, which have mutations in the RBD: Alpha/UK/B.1.1.7 (N501Y); Beta/South Africa/B.1.351 (K417N, E484K, and N501Y); Gamma/Brazil/P.1 (K417T, E484K, and N501Y); and Delta/India/B.1.617.2 (L452R and T478K)^29, 30^. The SARS-CoV-2 virus strains having mutations in the residues E484 or L452 frequently escape neutralizing antibodies^31–33^. We checked which clones could detect which spike variants via microscopy observation. Two clones (C17 and P17) exhibited severely reduced sensitivity for the spike variant carrying the E484K mutation; previously reported SARS72^34^, mNb6^35^, and Ty1^36^ nanobodies also did not detect these variants. Although the Ty1 nanobody did not sense the spike variant having the L452R mutation, the clones provided in this study remained able to detect the L452R substitution (Extended Data Fig. 4).

## Nanobodies for test kits

The finding that some clones recognized fluctuating, immature, or possibly unfolded SARS-CoV-2 spike proteins led us to consider whether the clones could detect the denatured SARS-CoV-2 spike variants via immunological assays. We sampled cell lysates expressing full-length C-terminally C9-tagged SARS-CoV-2 spike variants in Laemmli’s sodium dodecyl sulfate (SDS) buffer under reducing conditions^37^: The P158, P334, and P543 clones specifically detected SARS-CoV-2 spike proteins with the same sensitivity as the mouse monoclonal C9-tagged antibody (Fig. 2a). The banding patterns suggested that each clone recognized a part of the S1 domain. These clones (P158, P334, and P543) detected SARS-CoV-2 spike variants—including alpha and beta variants—via Western blotting.

**Fig. 2.**
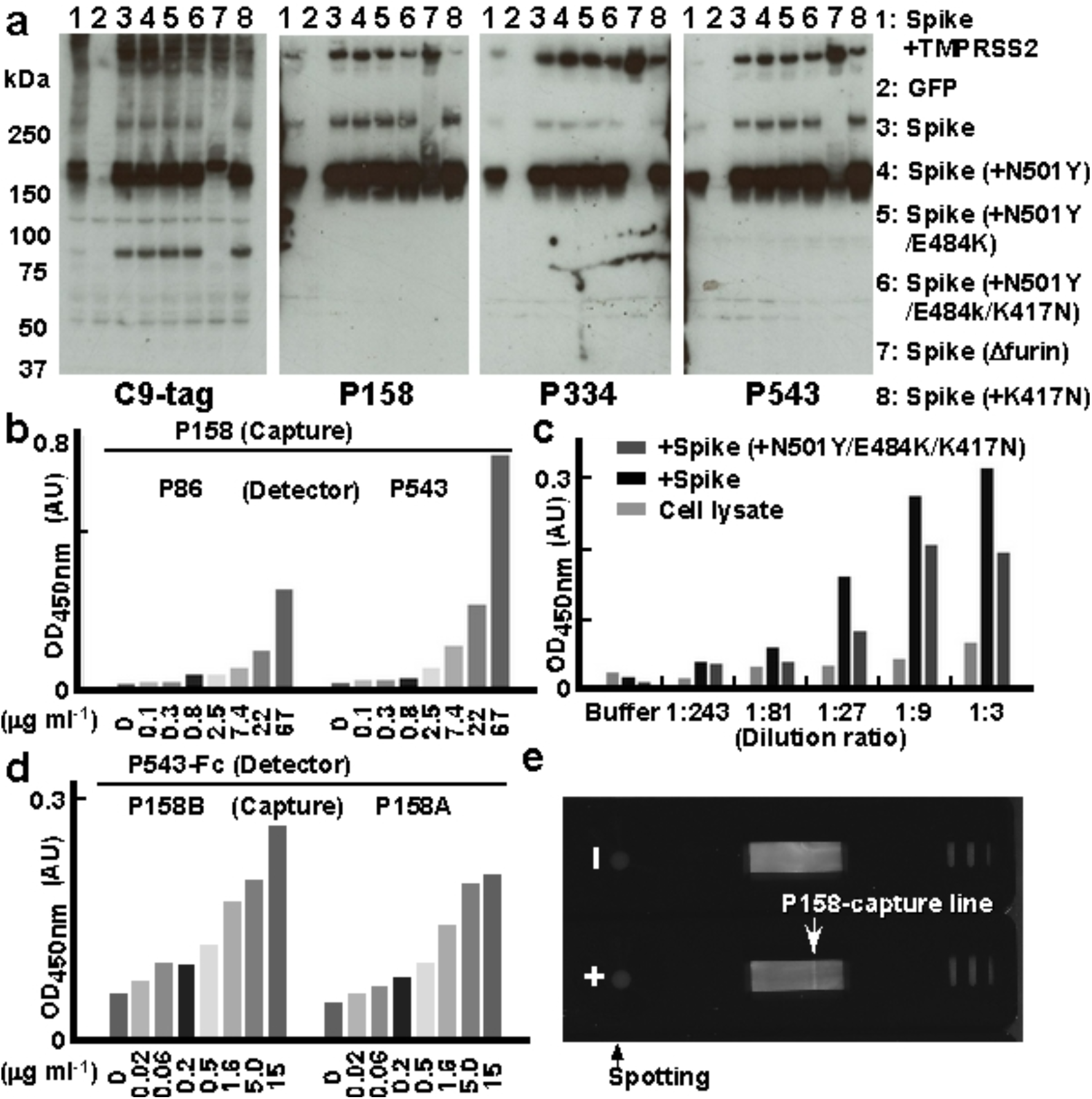
Nanobodies for Western blotting assays, ELISAs, and lateral flow assays. **a**, Nanobody-based Western blotting assays of SARS-CoV-2 spikes: lysates of HEK cells expressing GFP or C-terminally C9-tagged SARS-CoV-2 spike (D614G) variants (indicated with numbers) were blotted with the indicated nanobodies. **b-d**, Sandwich ELISA: **b**, signals of gradually diluted sample concentrations using P158 as the capture antibody and P86 or P543 (**b,c**) or P543-Fc (**d**) as the detection antibodies are graphed as a mean of the measurements from 2 wells (n=2). In (**c**), sequentially diluted lysates of HEK cells expressing the original SARS-CoV-2 spike (D614G) or beta variant (D614G, N501Y, E484K, and K417N) were assayed. **e**, A photograph of the results from lateral flow nitrocellulose membrane assays: P158 for the capture line (allow head) and P86 for the detection beads; 300 ng of the purified SARS-CoV-2 spike delta variant (D614G, L452R, and T478K) (+) or PBS (–) was spotted.

## Nanobody-based ELISA and lateral flow assay

We then tried to develop a sandwich ELISA kit to detect the SARS-CoV-2 spike using these nanobodies. We tested eight clones as detector nanobodies with six blocking conditions on P158-coated plates; we found that P86 and P543 worked well (Extended Data Fig. 5). The detection limit of the SARS-CoV-2 spike was approximately 1 μg ml^−1^ when using P158 as the capture and P543 as the detector nanobodies^38^ (Fig. 2b). Both P543 and P86 were able to specifically detect the SARS-CoV-2 spike original and beta variant in cellular homogenates: HEK cells expressing the full-length spikes were lysed and sequentially diluted (Fig. 2c). When the detection antibody was changed to Fc-tagged P543 (P543-Fc), the detection limit reached below 20 ng ml^−1^ (Fig. 2d). Moreover, antigen test kits based on a lateral flow membrane assay in which P158 lined a nitrocellulose membrane and P86 was conjugated to labelled beads were also able to detect 300 ng of the spike delta variant (Fig. 2e). We checked the capability of each clone via microscopy observations using cells expressing SARS-CoV-2 spike variants carrying associated mutations. The clones that were useful for blotting (P158, P334, and P543) and P86 were still able to detect the SARS-CoV-2 spike carrying mutations in the S1 domain, including deletion mutations (del) of H69, V70, Y144, Y145, L242, A243, and L244 and point mutations of L18F, T20N, P26S, D138Y, R190S, D215G, R246I, N439Y, Y453F, T478K, and A570D^39–42^ (Extended Data Fig. 6).

## Neutralizing activity of the nanobodies against SARS-CoV-2 spike variants

We assayed whether these clones could neutralize virus infection using SARS-CoV-2 spike pseudotyped HIV-1-based viruses. We used K562 cells stably expressing human ACE2 and serine protease TMPRSS2. We also produced pseudotyped viruses encoding the luciferase gene and expressing the SARS-CoV-2 spike variants. Five (C17, P17, P86, C116, and C246) out of nine clones inhibited the pseudotyped virus expressing the spike (original WH-1: D614G) in a dose-dependent manner (Fig. 3a). Then, we also tested the potency of the five clones for the other SARS-CoV-2 variants—alpha, beta, and delta. The P17, P86, and C116 clones more potently neutralized the alpha variant than Ty1; the P86 and C246 clones significantly suppressed the beta variant; and P17 and C246 suppressed the delta variant (Fig. 3a). The IC50 results were summarized and compared: C246, which stained intracellular spike (Extended Data Fig. 4c), suppressed all variants tested; P86 neutralized beta but not delta variants; and P17 neutralized delta but not beta variants (Fig. 3b). Because it was unclear to us why P17 could be affected by the E484K mutation (Beta) and P86 by the L452R mutation (Delta), we finally determined the differences in the epitopes of the two clones.

**Fig. 3.**
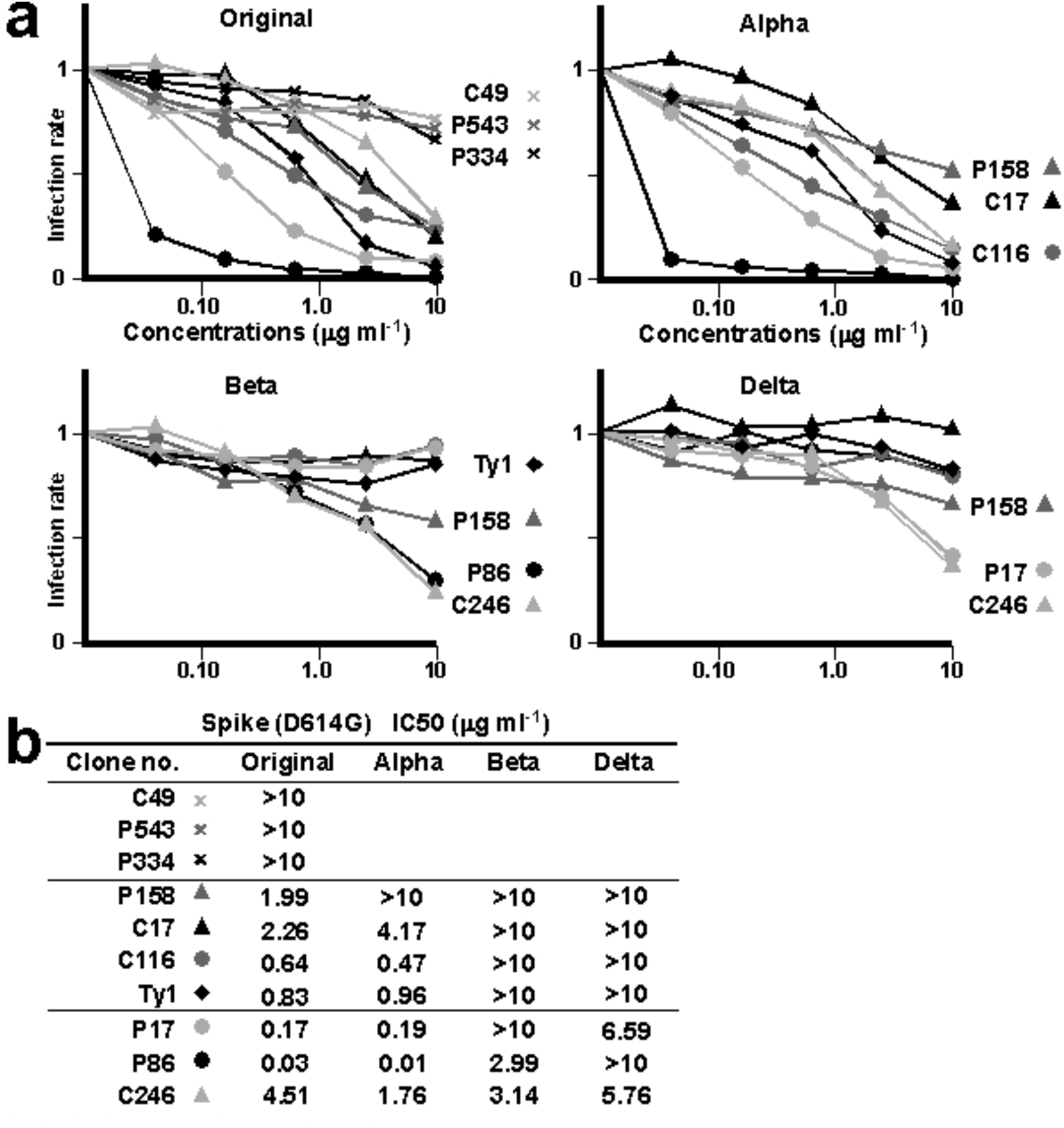
Pseudotyped virus neutralization assay. **a**, Means of infecting units of virus for the indicated SARS-CoV-2 spike variants are graphed (n=3). K562 cells expressing ACE2 and TMPRSS2 infected with HIV-1-based pseudotyped virus were assessed via luciferase assay. Relative infection units at the indicated concentrations of the nanobodies, which are marked individually, are plotted. **b,** Measured IC50s for each variant are summarized.

## The binding epitopes of the P86 and P17 clones

To determine the epitopes of the P86 and P17 clones on the SARS-CoV-2 spike, we performed a single-particle analysis of the spike-P86 and spike-P17 complexes using cryo-electron microscopy (cryo-EM). We observed 2-RBD-up (2-up+P86) and 3-RBD-up (3-up+P86) from the spike-P86 complex and 1-RBD-up (1-up+P17) and 2-RBD-up (2-up+P17) from the spike-P17 complex (Fig. 4a,b, Extended Data Fig. 7, and Extended Data Table 1). The densities of P86 and P17 were observed just the outside surfaces of both up-and down-RBDs—this arrangement was more apparent for the down-RBD (Extended Data Fig. 7). We observed that the densities of the clones bound to the down-RBD extended to the NTD of the neighbouring protomer (PDB entry: 7KRR)—more apparently in P86—the densities of P17 were too weak to determine the orientation (Fig. 4c,d).

**Fig. 4.**
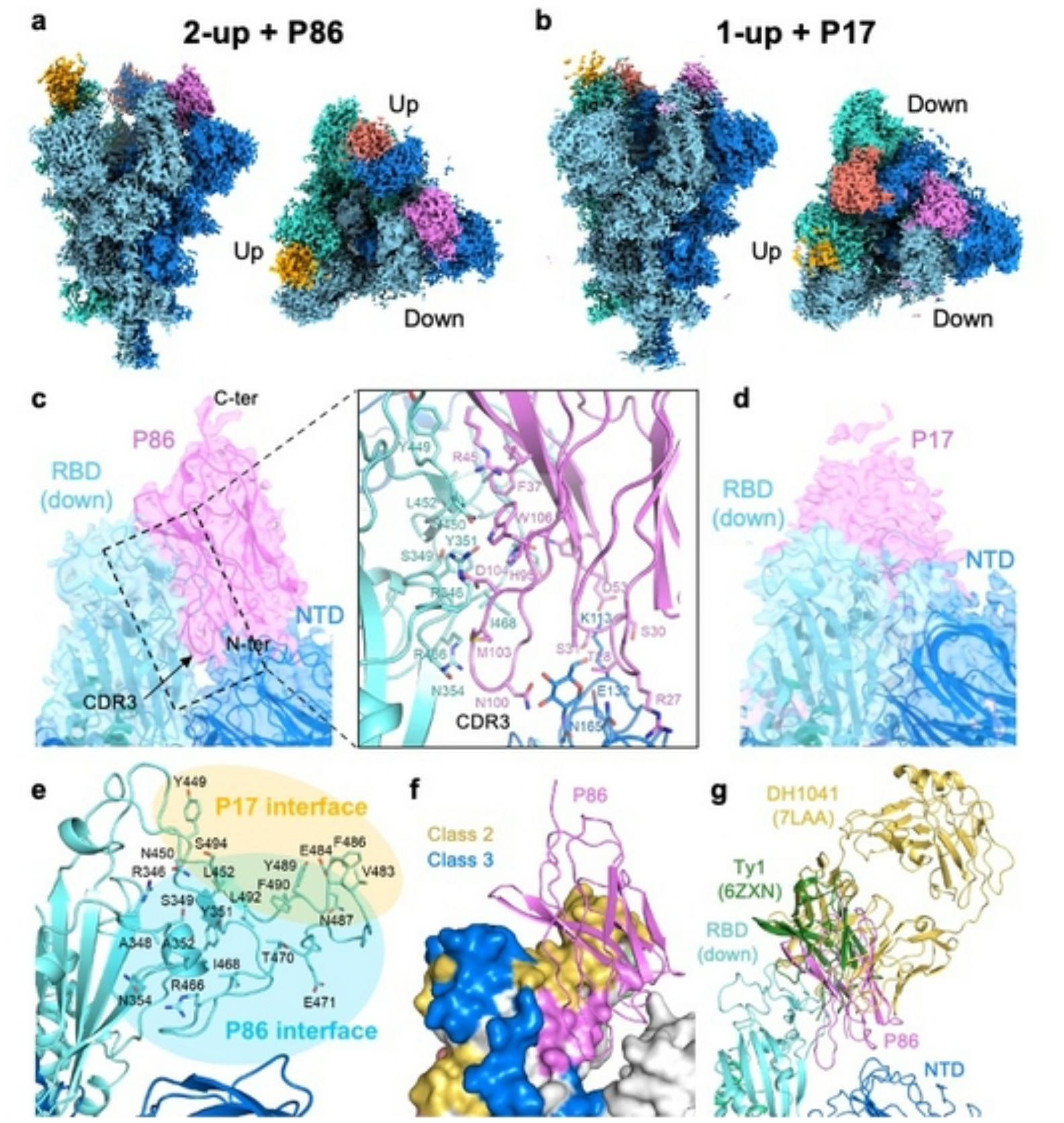
Cryo-EM density maps of SARS-CoV-2 spike-P86 and SARS-CoV-2 spike-P17 complexes. **a,b**, Final sharpened maps of the (**a**) 2-up+P86 and (**b**) 1-up+P17 datasets. The map regions corresponding to one protomer in the spike protein are coloured differently in cyan, blue, and turquoise. The map regions corresponding to one P86 or P17 molecule bound to the RBD regions are coloured magenta, red, and orange, respectively. **c**, Close-up view around P86 bound to the down-RBD with our fitted model. Right panel shows close-up view around the interface among P86, the down-RBD, and the NTD of the neighbouring protomer. **d**, Close-up view around P17 bound to the down-RBD. The model of D614G spike trimer (PDB entry: 7KRR) was fitted on the maps^50^. **e**, The residues located on the interface between the down-RBD and P86 or P17. **f**, Epitope mapping on the down-RBD. The epitopes of Class-2, Class-3, and P86 are shown in yellow, blue, and magenta, respectively. The colouring priority of the overlapping region is Class-3>Class-2>P86. The model of P86 is a ribbon structure. **g**, Structure comparison with other antibodies bound to the down-RBD. The structures are superposed on the single RBD region.

To build an atomic model of 2-up+P86, we determined the crystal structure of P86 at 1.60 Å resolution (Extended Data Table 2). The asymmetric unit contained two structurally identical monomers (a root mean square deviation for C_α_ atoms of 0.20 Å); the three CDR regions were visualized well (Extended Data Fig. 8a,b). We fit the P86 crystal structure to the density near the down-RBD in a locally refined map. The map showed two featured regions: a loop-like density between the down-RBD and the NTD of the neighbouring protomer and an elongated density located on the top (Extended Data Fig. 8c). We fit the CDR3 and the C-terminal (C-ter.) region into the former and the latter, respectively, and the whole body was adequately accommodated. After manual model modifications, the P86 monomer fit well into the final sharpened map (Fig. 4c [left panel] and Extended Data Fig. 8c-e). The edge of the CDR3 region was inserted into the cleft between the down-RBD and the NTD of the neighbouring protomer, forming many hydrophilic and hydrophobic interactions (Fig. 4c [right panel]). Compared to the density of P86, the density of P17 was shifted towards the top of the RBD (Fig. 4e). Therefore, the interface between the down-RBD and P17 was near the receptor-binding motif (RBM: ACE2-binding region); on the other hand, the interface between the down-RBD and P86 shifted to the NTD of the neighbouring protomer (Fig. 4e).

To date, human antibodies target four epitopes, all on the RBD: Class-1 (RBM class 1), Class-2 (RBM class 2), Class-3 (RBD core 1), and Class-4 (RBD core 2)^43–45^ (Extended Data Fig. 8f). The epitope of P86 was located outside of Class-2 and Class-3 and was on the opposite side of Class-1 and Class-4 (Fig. 4f and Extended Data Fig. 8f,g). We compared two antibodies—DH1041(PDB entry: 7LAA) and Ty1 (PDB entry: 6ZXN)—that target the near-surface of the down-RBD where P86 is bound^36, 46^. We superposed them on the single down-RBD within the 2-up+P86 model, which clearly showed P86—especially the CDR1 and CDR3 loops—uniquely bound the down-RBD and the NTD of the neighbouring protomer (Fig. 4c,g). We noticed that the P86-bound down-RBD interfered with the ACE2 to access the neighbouring up-RBD (Extended Data Fig. 8d). It seems that P86 is small enough to access the *hidden* cleft that is not recognized by human antibodies.

## Discussion

We provide the sequences of the nanobodies recognizing recently emerging SARS-CoV-2 spike VOCs. Only sequence information or expression vectors are sufficient to produce these clones in basic biological laboratories. We showed that the nanobodies detected the spike variants via ELISA, immunochromatography, and blotting assays. Therefore, these clones can be applied for surveillance of the virus in wastewater, monitoring infected individuals (e.g., passengers, farm animals, or pets), and self-testing. Here, we showed that C246, which recognized immature or unfolded spikes, reduced the activity of the three SARS-CoV-2 VOCs. We have not yet determined the epitope of C246: we wondered whether C246 might inhibit the maturation of the virus rather than compete for association with the target^47^.

The epitopes of P17 and P86 were shifted relative to each other (Fig. 4e), which caused different tolerances to the mutations—P17 neutralized the delta (L452R) variant, whereas P86 neutralized the beta (E484K) variant. The L452R variant may substantially reduce the avidity of P86 to the RBD: R452 is located near the centre of the P86 binding interface. By contrast, because P17 binds to the surface near the top of the RBD and R452 exists on the edge of the interface, P17 tolerates the long R452 side chain. E484 is part of the flexible *hook* rope of the RBD. Structural and molecular dynamics studies have suggested that the E484K mutation disorders the hook region^48^. We assume that the Class-2 antibodies can only bind a solid state of the hook containing E484, whereas P86 is not severely influenced by states of the hook region containing K484 because P86 also contacts the conserved cleft between the RBD and the NTD of the neighbouring protomer. The epitope of P86 seems too narrow to be accessed by conventional antibodies. Indeed, the region of the RBD where P86 binds has not yet been classified as an epitope of human antibodies^49^ (Extended Data Fig. 8f,g). Therefore, we expect these clones to be similar to *lime in a gimlet* of antibody-based therapies.

## Methods

### Ethics statement for animal care

Two young alpacas (*Vicugna pacos*) half-siblings—a 19-month-old male named “Puta” and a 19-month-old female named “Christy”—were immunized. Veterinarians of the KYODOKEN Institute for Animal Science Research and Development (Kyoto, Japan) bred, maintained health, recorded conditions, and performed the immunization studies by adhering to the published Guidelines for Proper Conduct of Animal Experiments by the Science Council of Japan. The KYODOKEN Institutional Animal Care and Use Committee approved the protocols for these studies (approval number 20200312) and monitored health conditions. The veterinarians immunized the alpacas with antigens and collected blood samples under anaesthesia.

### Immunization and library generation

Immunized antigens were purified recombinant CoV-2 spike complexes: the extracellular domain of the original WH-1 protein (GenBank: QHD43416) with or without the D614G mutation that carried a maintained or mutated furin cleavage site— N^679^SPRRA or IL^680^. The protein mixture emulsified in complete Freund’s adjuvant was subcutaneously injected into the two alpacas up to 9 times at 2-week intervals. Blood samples were collected from the jugular vein; peripheral blood mononuclear cells (PBMCs) were obtained with a sucrose density gradient using Ficoll^51^ (Nacalai Tesque, Kyoto, Japan). The PBMC samples were washed with PBS and suspended in RNAlater solution (Thermo Fisher Scientific K.K., Tokyo, Japan). Total RNA was isolated from the PBMC samples (Direct-Zol RNA MiniPrep: Zymo Research, Irvine, CA).

Complementary DNA was synthesized from 1 μg of the total RNA as a template with random hexamer primers and using SuperScript II reverse transcriptase (Thermo). Coding regions of the heavy-chain variable domains were amplified using LA Taq polymerase (TAKARA Bio Inc., Shiga, Japan) with two PAGE-purified primers (CALL001: 5’-GTCCTGGCTGCTCTTCTACAAGG-3’ and CALL002: 5’-GGTACGTGCTGTTGAACTGTTCC-3’). The amplified coding gene fragments of heavy-chain variable domains were separated on a 1.5% low-temperature melting agarose gel (Lonza Group AG, Basel, Switzerland). Approximately 700 base pair bands corresponding to the heavy-chain only immunoglobulin were extracted (QIAquick Gel Extraction Kit: Qiagen K.K., Tokyo, Japan). Nested PCR was performed to amplify coding genes of VHH domains using the VHH-PstI-For and VHH-BstEII-Rev primers and subcloned into the pMES4 phagemid vector^5, 52^ (GeneArt DNA Synthesis: Thermo). Electroporation-competent *Escherichia. coli* TG1 cells (Agilent Technologies Japan, Ltd., Tokyo, Japan) were transformed with the ligated plasmids under chilled conditions (Bio-Rad Laboratories, Inc., Hercules, CA). Colony-forming units of the libraries were checked with limiting dilution to maintain >10^7^ per microlitre. Colonies from 8 ml of cultured cells were harvested, pooled, and reserved in frozen glycerol stocks as parent libraries.

### Plasmid construction for protein expression

The gene encoding the extracellular domain of the SARS-CoV-2 spike protein (residues 31-1213) was codon optimized and synthesized into the pcDNA3.1(+) vector (Thermo) with an N-terminally modified IL-2-derived signal peptide (ILco2: MRRMQLLLLIALSLALVTNS)^53^; proline substitutions at residues K986P and V987P; and the C-terminal T4-phage fibritin trimerization domain (foldon) following a 6×His-tag^34, 54–56^. The expression vector of the SARS-CoV-2 S2 domain (residues 744-1213) was constructed by removing an N-terminal part of the extracellular domain of the SARS-CoV-2 spike (residues 31-743) and subcloned into the pcDNA3.1(+) vector. The expression vectors of the full-length SARS-CoV-2 spike and the human serine protease TMPRSS2 with a C-terminal C9-tag (TETSQVAPA) were acquired from AddGene (Summit Pharmaceutical International, Tokyo, Japan). The genes encoding the extracellular domain of human ACE2 (residues 1-614) and the endemic human coronavirus HCoV-OC43 (residues 1-1322) were codon optimized, synthesized with a C-terminal 6×His-tag and subcloned into the pcDNA3.1(+) vector. The whole gene of the human apolipoprotein B messenger-RNA-editing enzyme catalytic polypeptide-like (APOBEC) 3G (A3G)^26^ was also codon optimized, synthesized with a C-terminal 6×His-tag, and subcloned into the pcDNA3.1(+) vector.

For structural analyses, the sequence encoding the spike ectodomain (residues 1-1208) with proline substitutions, a “GSAS” substitution at the furin cleavage site (residues 682-685)^57^, and the C-terminal foldon trimerization motif followed by an 8×His-tag was cloned into the pcDNA3.1(+) expression vector (Invitrogen). Furthermore, the D614G mutation in the spike protein was introduced by the inverse PCR method. For cryo-EM, recombinant spike proteins were transiently expressed in Expi293-F cells (Thermo) maintained in HE400AZ medium (Gmep, Inc., Fukuoka, Japan). The expression vector was transfected using a Gxpress 293 Transfection Kit (Gmep) according to the manufacturer’s protocol. The culture supernatants were harvested five days post-transfection. The C-terminally 6×His-tagged spike proteins were purified using a nickel Sepharose 6 FF column (Cytiva) and size exclusion chromatography using a Superdex200increase 10/300 GL column (Cytiva) with buffer containing 50 mM HEPES (pH 7.0) and 200 mM NaCl.

### Biopanning

The purified proteins were i) the extracellular domain of the SARS CoV-2 spike; ii) only the S2 domain of the SARS CoV-2 spike; iii) the extracellular domain of the seasonal cold coronavirus spike of HCoV-OC43; iv) A3G; and v) homemade IgM coupled to N-hydroxysuccinimide (NHS)-activated magnet beads (Dynabeads, Thermo). One round of biopanning was performed using different protein-coated magnet beads in 50 mM phosphate buffer (pH 7.4) containing 1% (n-dodecyl-β-D-maltopyranoside) (DDM: Nacalai), 0.1% 3-[(3-cholamidopropyl)dimethylammonio]-1-propane sulfonate (CHAPS: Nacalai), 0.001% cholesterol hydrogen succuinate (CHS: Tokyo Chemical Industry Co., Ltd. (TCI), Tokyo, Japan); 0.1% LMNG (anatrace, Maumee, OH); and 500 mM NaCl. After 3 washes with the same buffer, the remaining phages bound to the washed beads were eluted with a trypsin-ethylenediaminetetraacetic acid (EDTA: Nacalai) solution at room temperature for 30 min. The elution was neutralized with a PBS-diluted protein inhibitor cocktail (c*O*mplete, EDTA-free, protease inhibitor cocktail tablets: Roche Diagnostics GmbH, Mannheim, Germany) and used to infect electroporation-competent cells. The infected cells were cultured and selected in LB broth Miller containing 100 μg ml^−1^ ampicillin (Nacalai) at 37°C overnight. The selected phagemids were collected using a QIAprep Miniprep Kit (Qiagen).

### Sequence analysis of nanobody libraries

The VHH-coding regions within parent libraries and 1-round target-enriched sublibraries were PCR amplified and purified using AMPure XP beads (Beckman Coulter, High Wycombe, UK). Then, dual-indexed libraries were prepared and sequenced on an Illumina MiSeq (Illumina, San Diego, CA) using a MiSeq Reagent Kit v3 with paired-end 300 bp reads^58^ (Bioengineering Lab. Co., Ltd., Kanagawa, Japan). Approximately 100,000 paired reads of each library were generated. The raw data of reads were trimmed of the adaptor sequence using cutadapt v1.18^59^, and low-quality reads were subsequently removed using Trimmomatic v0.39^60^. The remaining paired reads were merged using fastq-join^61^ and then translated to the amino acid sequences using EMBOSS v6.6.0.0^62^. Finally, unique amino acid sequences in each library were counted using a custom Python script combining seqkit v0.10.1^63^ and usearch v.11^64^. Enrichment scores of each clone were analysed by calculating the *P*-value of χ^2^ tests between the existing ratios among the different sublibraries. We chose nine clones whose enrichment scores in the SARS-CoV-2 spike biopanned sublibraries were higher than those in other biopanned sublibraries.

### Nanobody expression

Each selected amino-acid sequence was connected with a (GGGGS)_4_ linker as a tandem dimer; coding genes of these and of the previously reported SARS72 dimer, mNb6 dimer, and Ty1 monomer were codon-optimized and synthesized (Eurofins Genomics Inc, Tokyo, Japan). The synthesized genes were subcloned in the pMES4 vector to express N-terminal PelB signal peptide-conjugated and C-terminal 6×His-tagged nanobodies into the bacterial periplasm. These gene constructs were transformed into BL21(DE3) *E. coli* cells (BioDynamics Laboratory Inc., Tokyo, Japan) and plated on LB agar with ampicillin, which were incubated at 37°C overnight. Colonies were picked and cultured at 37°C to reach an OD of 0.6 AU; the cells were cultured at 37°C for 3 h or at 28°C overnight with 1 mM IPTG (isopropyl-β-D-thiogalactopyranoside: Nacalai). Cultured cells were collected by centrifugation. Nanobodies were eluted from the periplasm by soaking in a high osmotic buffer containing 200 mM Tris, 0.5 mM EDTA, and 500 mM sucrose (pH 8.0) at 4°C for 1 h. They were incubated with 2× volumes of a diluted buffer containing 50 mM Tris, 0.125 mM EDTA, and 125 mM sucrose (pH 8.0) with a trace amount of benzonase nuclease (Merck KGaA, Darmstadt, Germany) at 4°C for 45 min, and the supernatants were centrifuged (20,000×g, 4°C for 10 min). The supernatants were sterilized with the addition of gentamicin (Thermo) and passed through a 0.22-μm filter (Sartorius AG, Göttingen, Germany). The filtered supernatant was applied to a HisTrap HP nickel column (Cytiva) equipped on an ÄKTA purifier HPLC system (Cytiva) and washed; the bound His-tagged protein was eluted with 300 mM imidazole. The elution was concentrated with a VIVAspin 3,000-molecular weight cut off (MWCO) filter column (Sartorius) and applied to a Superdex75increase 10/300 GL gel-filtration column (Cytiva) equipped on an ÄKTA pure HPLC system (Cytiva) to obtain the dimer fractions and exclude cleaved monomers and imidazole. Purity was measured via Coomassie Brilliant Blue (CBB) staining.

### Antibodies

Antibodies used for Western blotting and cell staining were anti-His (rabbit polyclonal PM032: Medical and Biological Laboratories Co., Ltd. (MBL), Nagoya, Japan), anti-His (rabbit monoclonal EPR20547: ab213204, Abcam), and anti-rhodopsin C9 (TETSQVAPA) (mouse monoclonal 1D4, sc-57432: Santa Cruz Biotechnology Inc., CA). Horseradish peroxidase (HRP)-linked secondary antibodies included anti-mouse IgG (sheep polyclonal, NA931: GE Healthcare, Buckinghamshire, UK), anti-rabbit IgG (sheep polyclonal, NA934: GE Healthcare), and anti-alpaca IgG^65^ (goat polyclonal, 128-035-232: Jackson ImmunoResearch Laboratories, Inc., West Grove, PA), Alexa Fluor 488 goat anti-mouse IgG (rabbit polyclonal, P0449: Dako, Glostrup, Denmark), and Alexa Fluor 594 goat anti-rabbit IgG (Dako).

### Western blotting and CBB staining

Samples were incubated at 37°C for 30 min (for SARS-CoV-2 spike variants) or boiled for 2 min (for nanobodies) with Laemmli’s SDS sample buffer containing 2.5 mM Tris (pH 6.8), 2% SDS, 10% glycerol, 0.001% bromophenol blue, and 13.3 mM dithiothreitol (DTT). Samples were electrophoretically separated on 5–20% or 15–25% gradient polyacrylamide gels and electrophoretically transferred onto polyvinylidene difluoride (PVDF) membranes (Immobilon-P. Millipore, Billerica, MA). Blotted membranes were incubated overnight at 4°C with the C9 antibody or the C-terminally 6×His-tagged homodimer of nanobodies—the dilution ratios of P158, P334, and P543 were 1:5000, 1:1000, and 1:2500, respectively—in Tris-buffered saline (TBS, pH 7.4) containing 0.005% Tween 20 (TBST) and 5% skim milk. In the case of nanobody-based blotting, after 3 washes with TBST, the membranes were incubated with 1:5000-diluted anti-His-tag antibody (MBL) in TBST containing 5% skim milk at room temperature for 1 h. The membranes were soaked with 1:5000-diluted HRP-conjugated anti-rabbit or anti-mouse IgG secondary antibodies (GE Healthcare) in TBST containing 5% skim milk for 30 min at room temperature. After 3 washes with TBST, reactive protein bands were visualized using an ECL Plus system (Cytiva). For CBB R-250 staining, a Rapid Stain CBB Kit (Nacalai) was used according to the manufacturer’s protocol.

### Column chromatography for protein purification

HEK cells expressing the SARS-CoV-2 S2 domain or the extracellular domain of the HCoV-OC43 spike were cultured in serum-free Opti-MEM (modified Eagle’s medium: Thermo) containing 1% penicillin and streptomycin. After 48 h, the culture supernatants were centrifuged, filtered, and concentrated with VIVAspin 20 size exclusion columns (30,000-MWCO). Pellets of HEK cells expressing the extracellular domain of the SARS-CoV-2 spike were suspended in 50 mM phosphate buffer (pH 7.4) containing 1% DDM, 0.1% CHAPS, 0.001% CHS, 0.1% LMNG, and 500 mM NaCl. The suspension was passed through a 21-gauge needle (Terumo Co., Tokyo, Japan) several times, trace amounts of benzonase nuclease (Merck) were added, and the lysates were incubated at 4°C overnight. The cell lysate was cleared by centrifugation and filtration. The C-terminal 6×His-tagged spike protein was purified through a HisTrap HP 1 ml column equipped on an ÄKTA pure HPLC system (Cytiva) under step-by-step elution conditions using running (20 mM imidazole) and eluting (500 mM imidazole) phosphate buffers containing 20 mM phosphate, 500 mM NaCl, 0.5% DDM, 0.1% CHAPS, 0.001% CHS, and 0.1% LMNG (for spike proteins from the cell lysate) (pH 7.4). Elution fractions of 6×His-tagged proteins were identified via Western blotting. Anti-6×His-tagged antibody-positive fractions were gathered and concentrated via VIVAspin 6 size exclusion columns (30,000-MWCO) to reach a volume under 0.5 ml. Size exclusion chromatography experiments were performed on a Superose6increase 10/300 GL column (Cytiva) with an ÄKTA pure HPLC system (Cytiva). Sample volumes were approximately 0.5 ml; injected samples were separated in PBS or PBS-LMNG (for SARS-CoV-2 spike protein) in a chilled chamber with a flow rate of 0.1 ml min^−1^. Chromatograms were monitored at 280 nm using a UV spectrophotometer. Elution fractions were identified via Western blotting, gathered, and concentrated via VIVAspin 6 size exclusion columns (30,000-MWCO). Protein concentrations were measured with a NanoDrop 2000c spectrophotometer (1 mA.U. at 280 nm was equivalent to 1 mg ml^−1^) (Thermo).

### Cell culture and transfection

HEK and K562 cells were grown in Dulbecco’s modified Eagle’s medium (DMEM: Invitrogen) supplemented with 10% foetal bovine serum (FBS) and antibiotics (1% penicillin and streptomycin). The cells were cultured in a humidified incubator with 5% CO_2_ at 37°C. The human ACE2 and TMPRSS2 genes were transduced into K562 cells with a lentivirus. The lentiviral vector pWPI-IRES-Bla-Ak-ACE2-TMPRSS2 was acquired from AddGene (plasmid #154983).

### Measuring titres and nanobody-based sandwich ELISA

Two micrograms of the recombinant extracellular domain of the SARS-CoV-2 spike or 10 μg of the homodimer of nanobodies was diluted with 10 ml of 0.1 M carbohydrate buffer (pH 9.8). Each well of a MaxiSorp 96-well plate (Thermo) was coated with 100 μl of the diluent at 4°C overnight. To measure the titres of the immunized two alpacas, the wells were washed 3 times with high-salt PBS-LMNG (500 mM NaCl and 0.001% LMNG) and blocked with the high-salt PBS-LMNG containing 1% FBS at room temperature for 1 h. For screening conditions with the nanobody-based sandwich ELISA for the SARS-CoV-2 spike, the wells were washed with PBST (0.05% Tween 20, pH 7.4) and were blocked at room temperature for 1 h with five kinds of blocking solutions: 1×Carbo-Free blocking solution (Vector Laboratories, Inc., Burlingame, CA); 5% bovine serum albumin (BSA) (Sigma-Aldrich) in PBST; 5% skim milk in PBST; 3% casein (Merck) in 20 mM TBS (pH 11.0); or 5% polyvinylpyrrolidone (PVP) average molecular weight 10,000 (Sigma-Aldrich) in PBST. For measuring titres, 10 μl of a 0.1% dilution of serum from immunized alpacas in PBST was added to the well and incubated at room temperature for 1 h. After 3 washes with PBST, HRP-conjugated anti-alpaca VHH antibody (Jackson) at a dilution of 1:5000 was reacted at room temperature for 30 min. For screening of nanobody-based sandwich ELISA conditions, 100 μl of 1% (w/v) of the extracellular domain of SARS-CoV-2 spike (1 μg 100 μl^−1^) in high-salt PBS-LMNG was added to each well and captured at room temperature for 1 h. The well was washed 3 times with the high-salt PBS-LMNG; each 30 ng of biotin-conjugated nanobody in 100 μl (0.03% w/v) of PBS-LMNG was soaked at room temperature for 30 min. After 3 washes with PBS-LMNG, 100 μl of HRP-conjugated streptavidin (TCI) at a dilution of 1:5000 in PBS-LMNG was incubated at room temperature for 30 min.

The amounts of remaining HRP conjugates after 3 washes with the buffer were measured with the addition of 100 μl of 50 mM phosphate-citrate buffer (pH 5.0) containing 0.4 μg of *o*-phenylenediamine dihydrochloride (OPD) (Sigma-Aldrich) to develop at room temperature for 30 min, after which the reaction was stopped with the addition of 10 μl of 5 M sulfuric acid (H_2_SO_4_). Each well was read at an optical density (OD) of 450 nm using a microplate reader (Bio-Rad).

### Immunochromatography

Antigen test kits detecting the SARS-CoV-2 spike based on nitrocellulose lateral flow assays were developed by Yamato Scientific Co., Ltd. (Tokyo, Japan). Briefly, purified P158 (330 ng μl^−1^) was lined approximately 1 mm wide on an IAB90 nitrocellulose membrane (Advantech, Toyo Roshi Kaisha, Ltd., Tokyo, Japan). The membrane was soaked in 1×Carbo-Free blocking solution at room temperature for 1 h and air dried. Purified P86 or P543 nanobodies were amine coupled to 30-nm diameter Estapore beads (Sigma-Aldrich) according to the manufacturer’s protocol. Glass fibre paper (GF/DVA: Cytiva) was soaked in blocking solution containing 0.005% (w/v) nanobody-coupled Estapore beads for saturation and then dried under vacuum conditions. The lined nitrocellulose membrane and the bead-absorbed glass fibre paper were overlaid on a backing sheet (Cytiva). CF4 paper (Cytiva) was used for both the sample pad and the absorbance pad; they were overlaid on the prepared backing sheet as well. The four-layered sheet was cut 5 mm wide and housed in black cases: the cassettes were stored in sealed packages with silica gels. When dried, 150 μl of sample was spotted onto the sample pad; the kits were photographed under a 315 nm UV lamp for an arbitrary amount of time.

### Kinetic assays via biolayer interferometry (BLI)

Real-time binding experiments were performed using an Octet Red96 instrument (fortèBIO, Pall Life Science, Portsmouth, NH). Each purified nanobody clone was biotinylated with EZ-Link Sulfo-NHS-LC-Biotin (Thermo) according to the manufacturer’s protocol; uncoupled biotin was excluded with a size exclusion spin column (PD SpinTrap G-25: Cytiva) in PBS (pH 7.4). Assays were performed at 30°C with shaking at 1,000 rpm. Biotin-conjugated clones at 10 μg ml^−1^ were captured on a streptavidin-coated sensor chip (SA: fortèBIO) to reach the signals at 4 nm. One uncoated sensor chip was monitored as the baseline; another biotin-conjugated anti-IL-6-coated sensor chip (anti-IL-6 nanobody: COGNANO Inc.) was monitored as the background. The remaining 6 channels were immobilized with biotinylated anti-SARS-CoV-2 spike clones, and real-time binding kinetics to the purified extracellular domain of the SARS-CoV-2 spike trimer complex were measured in sequentially diluted concentrations at the same time (8 channels per assay). The concentrations of the SARS-CoV-2 spike loaded varied between 1-32 μg ml^−1^, corresponding to 1-32 nM or less, when an average of the molecular weight of the SARS-CoV-2 spike trimer complex was estimated to be approximately 1,000 kDa or more according to chromatograms of gel filtration column chromatography (Extended Data Fig. 1b). Assays were performed with high-salt phosphate buffer containing 500 mM NaCl and 0.001% LMNG (stabilized spike: for P17, P86, P334, and C116) or with hypotonic phosphate buffer containing 25 mM NaCl and 0.00005% LMNG (fluctuated spike: for P158, P543, C17, C49, and C246). After baseline equilibration for 180 s in each buffer, the association and dissociation were carried out for 600 s each. The data were double subtracted before fitting was performed with a 1:1 fitting model in fortèBIO data analysis software. The equilibrium dissociation constant (*K*_D_), *k_dis_*, and *k_on_* values were determined with a global fit applied to all data.

### Pseudotyped virus production

HIV-1-based SARS-CoV-2 spike pseudotyped virus was prepared as follows: LentiX-HEK293T cells were transfected using a polyethyleneimine transfection reagent (Cytiva) with plasmids encoding the C-terminally C9-tagged full-length SARS-CoV-2 spike variants (original, alpha, beta, and delta) and HIV-1 transfer vector encoding a luciferase reporter, according to the manufacturer’s protocol. The mutations of each variant are as follows: original (D614G); alpha (H69del, V70del, Y144del, N501Y, A570D, D614G, P681H, T716I, S982A, and D1118H); beta (L18F, D80A, D215G, R246I, K417N, E484K, N501Y, D614G, and A701Y); and delta (T19R, G142D, E156del, F157del, R158G, L452R, T478K, D614G, P681R, and D950N). Cells were incubated for 4-6 h at 37°C with medium that was then replaced with DMEM containing 10% FBS for the following 48-h culture. The supernatants were then harvested, filtered through a 0.45-μm membrane, concentrated with ultracentrifugation, and frozen at –80°C.

### Pseudotyped virus neutralization assay

Fivefold sequentially diluted nanobodies were incubated with K562 cells expressing human ACE2 and TMPRSS2 for 1 h at 37°C. K562 cells were subsequently infected with SARS-CoV-2 pseudotyped viruses and cultured for 2 days. The cells were lysed, and luciferase activity was measured using the Steady-Glo Luciferase Assay System (Promega KK, Osaka, Japan) with a microplate spectrophotometer (ARVO X3: PerkinElmer Japan Co., Ltd., Kanagawa, Japan). The obtained relative luminescence units were normalized to those derived from cells infected with the SARS-CoV-2 pseudotyped virus in the absence of nanobodies.

### Flow cytometry

The ability of nanobodies to bind to the cell surface of the SARS-CoV-2 spike was studied by fluorescence-activated cell sorting (FACS)^66^. K562 cells expressing the SARS-CoV-2 spike were incubated with 1 μg ml^−1^ purified nanobody on ice for 30 min. After washing, the cells were incubated with an anti-His antibody (Abcam) on ice for 30 min and then Alexa 647-conjugated anti-rabbit IgG (Dako). The cells were analysed with a Beckman-Coulter FC-500 Analyzer (Coulter Electronics, Hialeah, FL). The Ty1 and B9^67^ nanobodies were used as controls. A region of positive signals for Ty1 was square-gated.

### Microscopy analyses for cell staining

HEK cells were transiently transfected with plasmids encoding the C-terminally C9-tagged full-length SARS-CoV-2 spike variants using Lipofectamine 3000 (Thermo) according to the manufacturer’s instructions. The next day, the cells were seeded on collagen type I-coated culture plates (IWAKI, AGC TECHNO GLASS CO., LTD., Shizuoka, Japan) and cultured for 24 h before being fixed with 2% paraformaldehyde (PFA) at 4°C overnight. After 3 washes with PBST (0.005% Tween), the cells were blocked with PBST containing 2% goat serum (blocking solution) at room temperature for 1 h. Each well was soaked with 100 μl of the blocking solution containing 30 ng of purified nanobody, except for 6 ng of C116, at 4°C overnight. After washing with PBST, an appropriately diluted anti-His-tagged antibody and anti-C9-tagged antibody in blocking buffer were added and reacted at room temperature for 1 h. Finally, after washing, fluorescently conjugated anti-rabbit IgG (594 nm emission) and anti-mouse IgG (488 nm emission) antibodies (Alexa Fluor: Thermo) were diluted with blocking buffer and added to the wells, and the fixed cells were labelled at room temperature for 1 h before washing 3 times with PBST. Cellular nuclei were visualized with 4’,6-diamidino-2-phenylindole (DAPI). Stained cells were imaged with a 2-ms exposure time (594 nm emission), with a 10-ms exposure time (488 nm emission), or automatically adjusted exposure time (DAPI) using microscopy (IX71S1F-3: Olympus Corporation, Tokyo, Japan) with the cellSens Standard 1.11 application (Olympus). A 3×4 cm^2^ printed rectangle corresponds to a 165×220 μm^2^ observed field.

### Expression and purification of monomeric P86 and P17

The monomeric C-terminally 6×His tagged nanobody genes were cloned into the pMES4 vector. The complete amino acid sequences are as follows (signal peptide sequence is underlined). P86: <u>MKYLLPTAAAGLLLLAAQPAMA</u>QVQLQESGGGLVQAGGSLR LSCVASGRTFSSLNIVWFRQAPGKERKFVAAINDRNTAYAESVKGRFTISRDNAK NTVHLQMNSLKPEDTAVYYCHSADVNGGMDYWGKGTQVTVSSHHHHHH. P17: <u>MKYLLPTAAAGLLLLAAQPAMA</u>QVQLQESGGGLVQAGGSLRLSCAASGR TSSVYNMAWFRQTPGKEREFVAAITGNGGTTLYADSVKGRLTISRGNAKNTVSLQMNVLKPDDTAVYYCAAGGWGKERNYAYWGQGTQVTVSSHHHHHH. Bacterial BL21(DE3) *E. coli* cells were transformed with the plasmids and grown on an LB ampicillin-supplemented plate; colonies were picked and inoculated into 5 ml of LB medium containing 200 μg ml^−1^ ampicillin; and the cells were cultured in a shaking incubator overnight at 37°C. The cultures were transferred to 1 L of LB medium containing 200 μg ml^−1^ ampicillin. At an optical density below 0.6, cells were cultured for 3 h at 37°C with 1 mM IPTG.

The bacterial pellet was collected by centrifuging at 6,000×g for 30 min and suspended in 45 ml of lysis buffer containing 20 mM Tris (pH 8.0), 0.68 mM EDTA, 500 mM sucrose, and a trace amount of benzonase nuclease (Merck). The solutions were mixed on a TR-118 tube rotator (AS ONE Corporation, Osaka, Japan); 90 mL of 20 mM Tris (pH 8.0) was added, and the mixtures were rotated for 45 min at 4°C. The resulting solutions were centrifuged at 20,000×g for 10 min at 4°C. The supernatant was filtered and loaded onto a Ni-NTA column (Cytiva) equilibrated with 20 mM Tris buffer (pH 8.0). The column was washed several times with 20 mM Tris buffer (pH 8.0) containing 40 mM imidazole for P86 or 20 mM Tris buffer (pH 8.0) containing 150 mM NaCl and 40 mM imidazole for P17. Then, the nanobody was eluted with 20 mM Tris buffer (pH 8.0) containing 250 mM imidazole for P86 or 150 mM NaCl and 350 mM imidazole for P17.

The fractions containing nanobodies were collected and further purified using a HiLoad 16/60 Superdex 75 gel filtration column (Cytiva) equilibrated with 20 mM Tris buffer (pH 8.0) containing 150 mM NaCl for P86 or with 20 mM HEPES buffer (pH 8.0) containing 150 mM NaCl for P17. The peak fractions were collected and concentrated. The final concentrations of the P86 and P17 sample were 51 mg ml^−1^ and 3.0 mg ml^−1^, respectively, based on the use of the absorption coefficient at 280 nm.

### Cryo-EM specimen preparation and data collection

An epoxidized graphene grid (EG-grid) was used to increase the number of protein particles. The trimer complex of the SARS-CoV-2 spike (D614G) with the furin-resistant mutation (“GSAS”) at a concentration of 0.1 mg ml^−1^ was mixed with a 5-time molar excess of P86 or P17 monomer and incubated on ice for 10 min, and 3 μl of the spike-nanobody complex solution was applied to the EG-grid. After incubation at room temperature for 5 min, the grids were blotted with a force of –3 and a time of 2 s in a Vitrobot Mark IV chamber (Thermo) equilibrated at 4°C and 100% humidity and then immediately plunged into liquid ethane. Excessive ethane was removed with filter paper, and the grids were stored in liquid nitrogen. All cryo-EM image datasets were acquired using a JEM-Z300FSC (CRYO ARM 300: JEOL, Tokyo, Japan) operated at 300 kV with a K3 direct electron detector (Gatan, Inc., Pleasanton, CA) in CDS mode^68^. The Ω-type in-column energy filter was operated with a slit width of 20 eV for zero-loss imaging. The nominal magnification was 60,000×, corresponding to 0.870 Å per pixel. Defocus varied between –0.5 μm and –2.0 μm. Each movie was fractionated into 60 frames (0.0505 s each, total exposure: 3.04 s) with a total dose of 60 e^−^/Å^2^.

### Cryo-EM image processing and refinement

The images were processed using RELION 3.1^69^. Movies were motion corrected using MotionCor2^70^, and the contrast transfer functions (CTFs) were estimated using CTFFIND 4.1^71^. Micrographs whose CTF max resolutions were beyond 5 Å were selected. Three-dimensional (3D) template-based autopicking was performed for all images, and the particles were extracted with 4× binning, which were subjected to two rounds of 2D classification. An initial model was generated and used as a reference for the following 3D classification in the P86 dataset. In the P17 dataset, a density map of spike trimers (our previous dataset) was used as a reference. Reference-based 3D classification (into 4 classes) without applying symmetry was conducted, and the selected particles were re-extracted without binning. 3D autorefinement without applying symmetry, soft mask generation, postprocessing, CTF refinement, Bayesian polishing, and another round of 3D autorefinement were performed. Then, focused 3D classification without alignment was performed to separate up-and down-states in one RBD. The selected particles were subjected to another round of 3D autorefinement, postprocessing, CTF refinement, 3D autorefinement. C3 symmetry was applied during the final round of 3D autorefinement for the 3-up+P86 dataset. The data were imported and further processed with non-uniform refinement in cryoSPARC v3.2.0^72^. The final map resolutions (FSC=0.143) were 3.03 Å and 2.70 Å in the 2-up and 3-up states in the P86 dataset and 3.20 Å and 3.29 Å in the 1-up and 2-up states in the P17 dataset, respectively. For the 2-up+P86 dataset, we tried local refinement to visualize the density of P86 bound to the down-RBD, but the resolution was limited to 5.13 Å (FSC=0.143).

The model was built for the 2-up+P86 dataset. The SARS-CoV-2 spike trimer D614G mutant (PDB entry: 7KRR)^50^ and the crystal structure of P86 were used initial models. After the models were manually fitted into the density using UCSF Chimera v1.15^73^ and modified in Coot v0.8.9.2^74^, real space refinement was performed in PHENIX v1.19.1^75^. The model was validated using MolProbity^76^, and this cycle was repeated several times. Figures were prepared using UCSF Chimera^73^, ChimeraX^77^, and PyMOL v2.5.0 (Schrödinger, LLC, New York, NY). The parameters are summarized in Extended Data Table 1.

### Crystallization of P86

Crystallization was performed by the sitting drop vapour diffusion method. A mosquito crystallization machine (TTP LabTech Ltd., Melbourn, Hertfordshire, UK) was used to prepare drops on 96-well VIOLAMO plates (AS ONE). The reservoir solution was 60 μl in volume, and 0.1 μl of protein solution was mixed with 0.1 μl of reservoir solution. Crystals appeared under the condition of 51 mg ml^−1^ protein, 0.2 M ammonium sulfate, and 30% (w/v) polyethylene glycol 4,000 at 20°C. A crystal cluster was crushed, and a peeled single crystal was harvested by LithoLoop (Protein Wave, Nara, Japan). Before the crystal was frozen in liquid nitrogen, it was soaked in the crystallization solution supplemented with 5% (v/v) ethylene glycol.

### X-ray data collection, processing, structure solution, and refinement

An X-ray diffraction experiment was performed on the BL44XU beamline of SPring-8 (Hyogo, Japan). Diffraction images were collected at 100 K using an EIGER X 16M detector (Dectris, Philadelphia, PA, USA). The beam size was 50.0×50.0 μm^2^ (h×w). A 0.8-mm Al attenuator was used to weaken the X-ray. The crystal-to-detector distance was 160 mm. The exposure time per frame and the oscillation angle were 0.1 s and 0.1°, respectively. A total of 1,800 images were collected. The dataset was processed using XDS^78^ and scaled by Aimless^79^. Molecular replacement phase determination was performed by MOLREP^80^ with a nanobody structure (PDB code ID: 5IVO)^81^ as a search model. Initial model building was performed by PhenixAutoBuild implemented in PHENIX^75^. Manual model building was performed using Coot^74^. The program refmac5^82^ in the ccp4 suite^83^ and the program Phenix-refine^75^ were used for structural refinement. The stereochemical quality of the final model was checked by Molprobity^76^. Data collection and refinement statistics are summarized in Extended Data Table 2.

#### Material availability

COGNANO Inc. will agree to the use of any materials and methods except for therapeutic use as long as appropriately referring to this paper or the company name; materials will be delivered through NITTOBO MEDICAL Corporation Ltd., Tokyo, Japan; and lateral flow assay (antigen test) kits will be developed and delivered through Yamato Scientific, Co., Ltd., Tokyo, Japan. Responsible requests for materials for research use should be directed to A.T.-K.

### Data availability

Density maps are available at the Electron microscopy Data Bank (EMDB) and Protein Data Bank (PDB) with accession codes EMD-32078 (2-up+P86), EMD-32079 (3-up+P86), EMD-32080 (1-up+P17), EMD-32081 (2-up+P17) and PDB-7VQ0 (2-up+P86). Additional cryo-EM data supporting this study are available from Ke.N. on reasonable request. The coordinate and structure factor files are deposited at the PDB (PDB code ID: 7VPY). Raw data are available at Integrated Resource for Reproducibility in Macromolecular Crystallography (https://proteindiffraction.org/).

## Acknowledgments

This study was supported by donations from POPURI Pharmacy Co., Ltd. (Kyoto, Japan), grants from the AMED Research Program on Emerging and Re-emerging Infectious Diseases (JP20fk0108268, JP20fk018517, and JP20fk018413 to A.T.-K.), JSPS KAKENHI (JP20K22630 to J.F. and JP25000013 to Ke.N.), Platform Project for Supporting Drug Discovery and Life Science Research (BINDS) from AMED (JP21am0101117 to Ke.N.), Cyclic Innovation for Clinical Empowerment (CiCLE) from AMED (JP17pc0101020 to Ke.N.), Program on Open Innovation Platform with Enterprises, Research Institute and Academia, Japan Science and Technology Agency (JST, OPERA, JPMJOP1861 to T.I.), JEOL YOKOGUSHI Research Alliance Laboratory of Osaka University to Ke.N, and KYOTO industrial Support Organization 21, the subsidies (Sangakukou no Mori) to COGNANO Inc. We thank all staff of Kyoto University Hospital, Kyoto University Medical Research Support Centre of Graduate School of Medicine, especially Yohta Fukuda, Haruyasu Asahara, and Maiko Moriguchi, Osaka University; the BL44XU beamline of SPring-8; and especially Kaori Yurugi, COGNANO Inc. We are pleased to thank Dr. Nakanishi and the veterinarians of KYODOKEN Institute for caring for animals and performing experiments. We also thank the Kyoto University Livestock Farm, which partially maintained the alpacas, and Satoshi Endo, the mayor of Hirono-machi town, Fukushima, and their local government, who granted construction of a farm to maintain alpacas.

## Author contributions

A.I. and A.T.-K. conceived the study. R.M. performed biochemical experiments, generated large libraries, performed biopanning, and assembled microscopy data. H.Y. performed bioinformatic analyses. Yo.K. and Ya.K. performed virus assays. K.K., Ka.N., Y.Y., K.M., A.R., K.Y., and I.A. prepared and certified materials. J.F. prepared cryo-EM specimens and collected data. J.F. and F.M. processed the cryo-EM images. J.F., T.I., and K.N. interpreted structures. K.Y. performed crystallographic studies. R.M., J.F., A.I., Ke.N., and A.T.-K analysed the data. R.M., J.F., Yo.K., Ya.K., K.K., H.Y., and K.Y. drew figures. R.M., J.F., K.Y., I.A., and A.T.-K. wrote the first draft of the manuscript. J.F., K.S., Y.M., T.I., A.I., Ke.N., and A.T.-K. reviewed and commented on the manuscript. All authors approved the reviewed manuscript.

---

**Extended Data Fig. 1.**
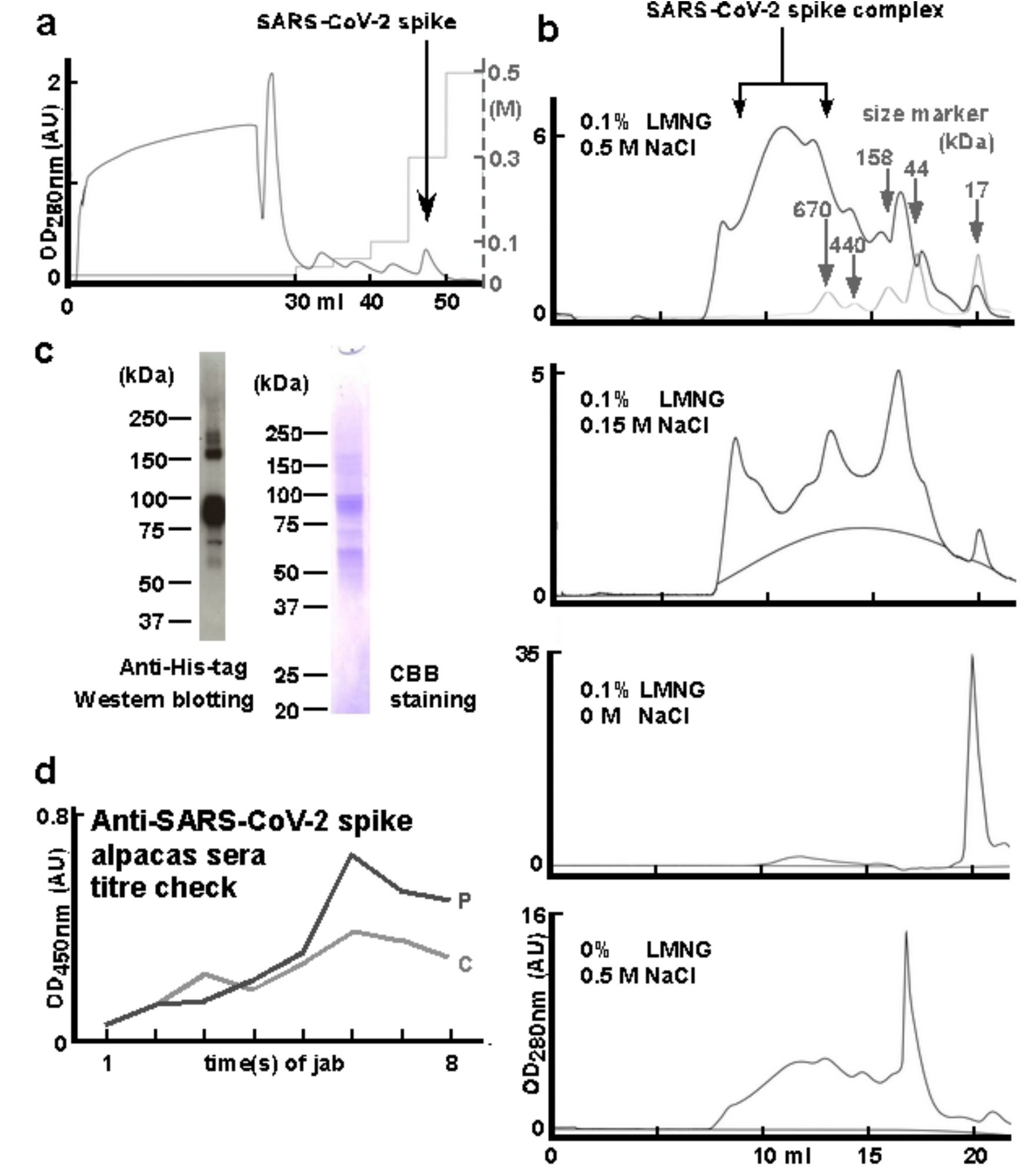
Purification of the extracellular SARS-CoV-2 spike complex. **a**, A chromatogram of the first purification step of the SARS-CoV-2 spike from HEK cell lysate using a nickel column: absorbance units (AU) of UV_280 nm_, amounts (ml) of buffer, concentrations (M) of imidazole, and the elution peak containing the SARS-CoV-2 spike. **b**, Chromatograms of the size-exclusion gel filtration of the recombinant SARS-CoV-2 spike complex under different conditions (concentrations of NaCl and LMNG). The first chromatogram is overlayed with those for marker proteins (arrowed). Two arrows indicate the elution peak of the SARS-CoV-2 spike. **c**, Western blotting analysis of the purified SARS-CoV-2 spike using an anti-His antibody and Coomassie Brilliant Blue (CBB) staining. **d**, Serum titres of the immunized alpacas—named P: Puta and C: Christy—for the SARS-CoV-2 spike were measured via ELISA and graphed: mean of the measurements for 2 wells.

**Extended Data Fig. 2.**
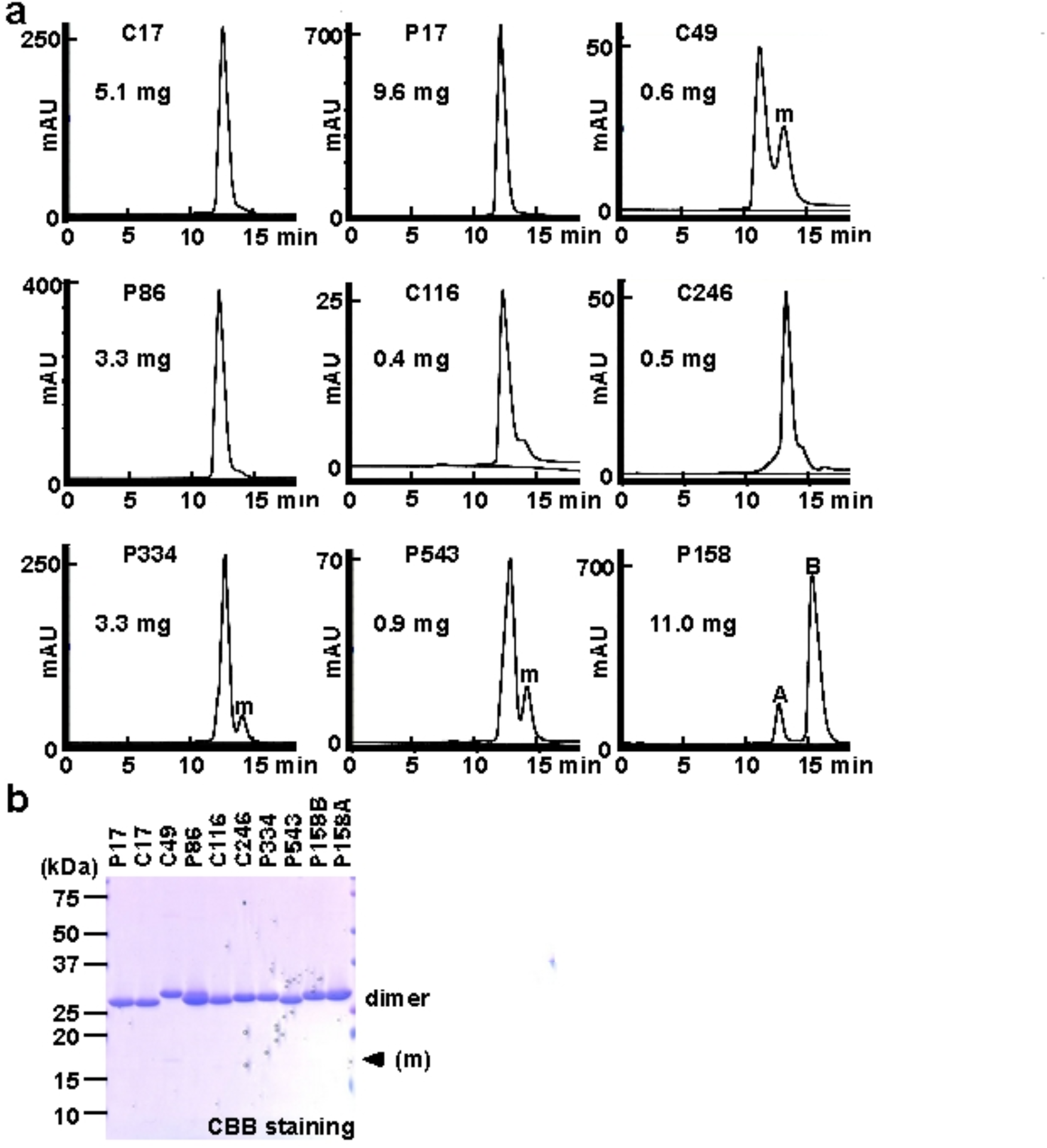
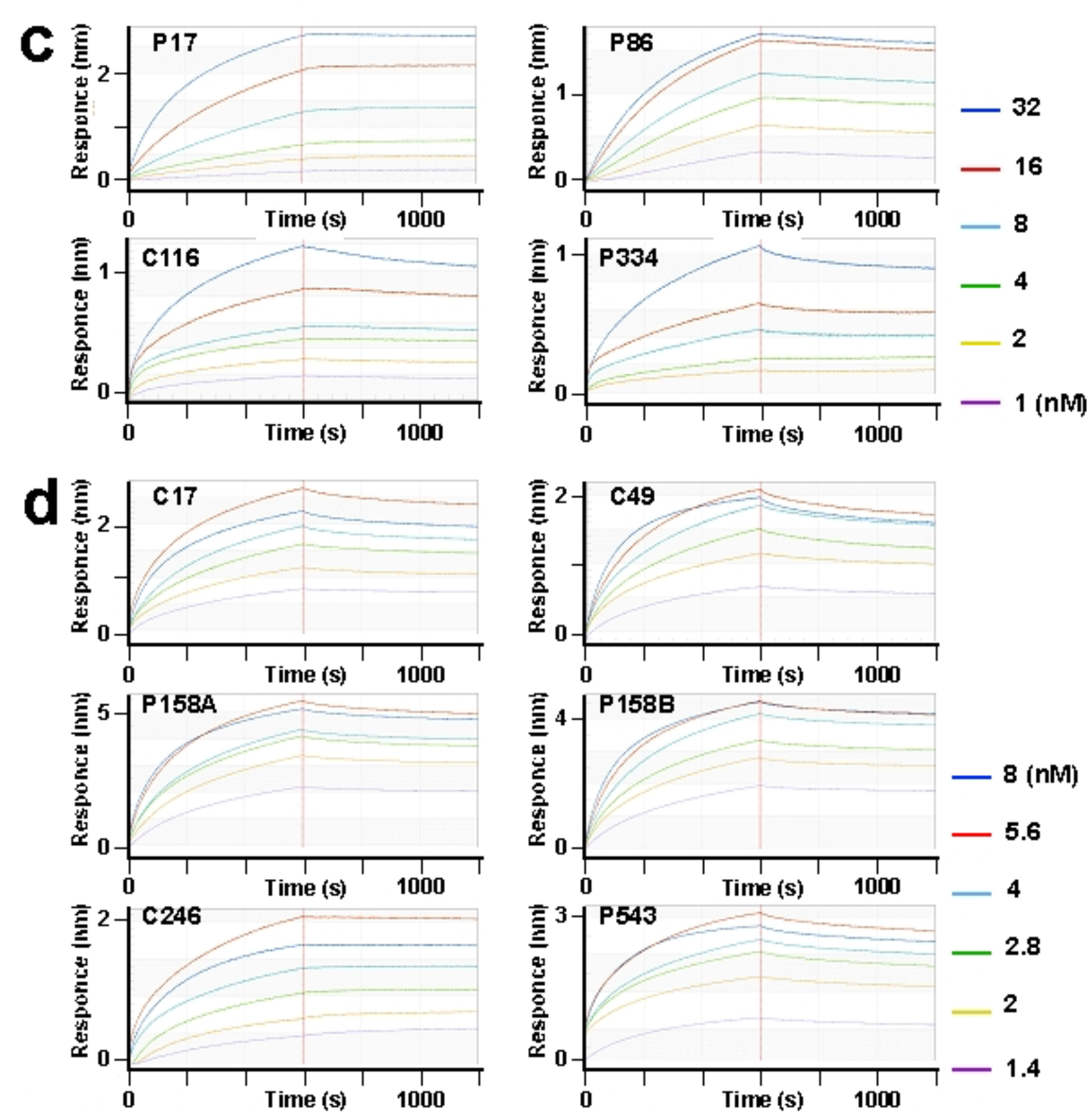
Purification and kinetic assays. **a**, Chromatograms showing the final purification step of the indicated clones: subpeaks confirmed as cleaved monomers are marked (m); the amounts of purified clones in dimers from 1 L-culture of BL21 bacteria are shown in insets. The dimer peaks observed for P158 are named A and B. **b**, The purified clones were analysed using CBB staining (m; cleaved monomer). Both the A and B peaks of P158 showed no differences via SDS-PAGE analysis. **c, d**, Sensorgrams of real-time kinetics assays using biolayer interferometry (BLI), showing five different concentrations of the purified SARS-CoV-2 spike in a high-salt buffer (**c**) or an anionic buffer (**d**).

**Extended Data Fig. 3.**
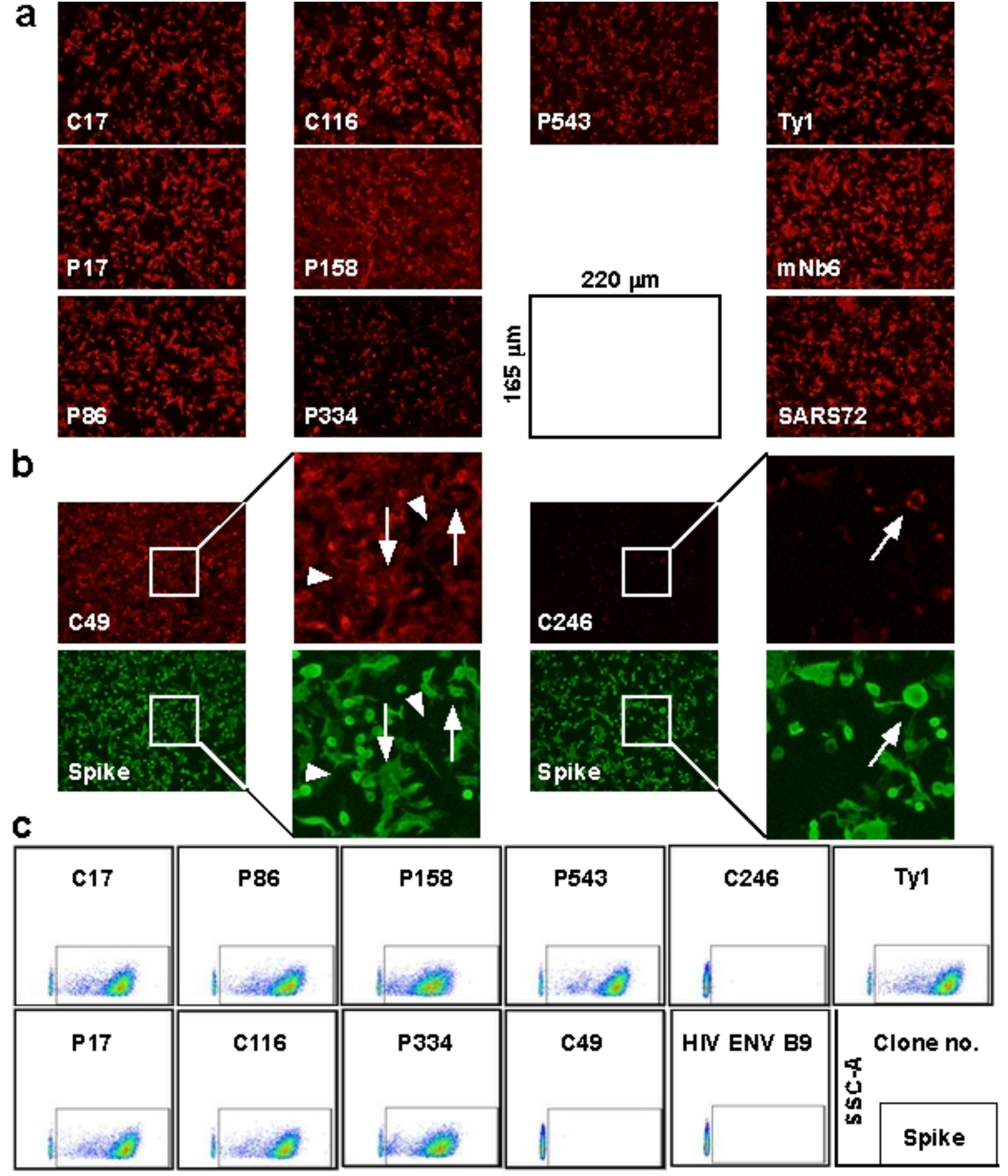
Microscopy and flow cytometric analyses of anti-spike nanobodies. **a,** Photographs of stained HEK cells expressing the C9-tagged SARS-CoV-2 spike. **b**, White squared regions show zoomed-in views: arrows indicate intracellularly stained SARS-CoV-2 spike (C49 and C246), and arrowheads indicate example background signals (C49). **c**, Flow cytometric analyses of K562 cells expressing the original SARS-CoV-2 spike—signals with Ty1 squared and B9: an anti-HIV-1 nanobody.

**Extended Data Fig. 4.**
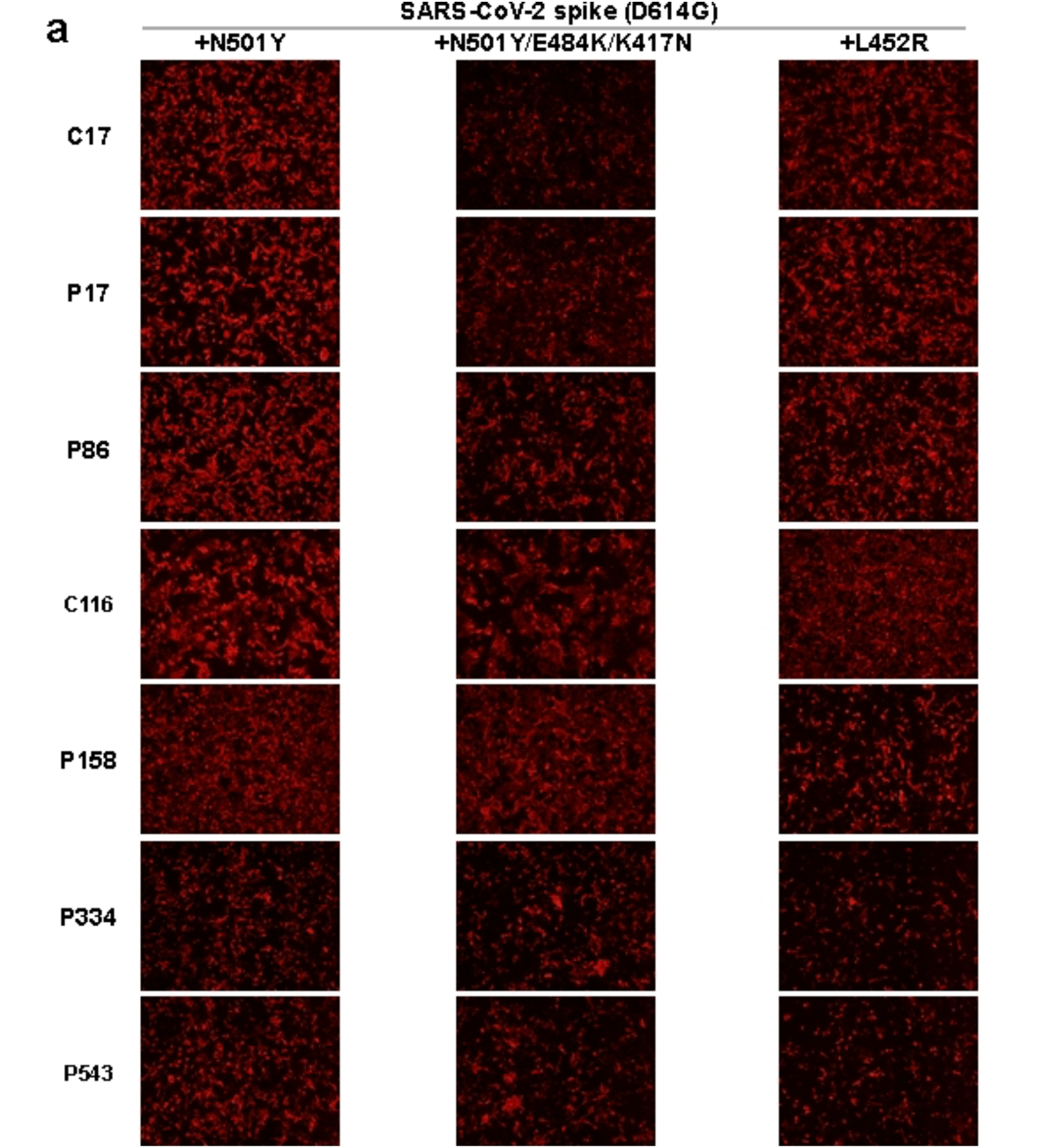
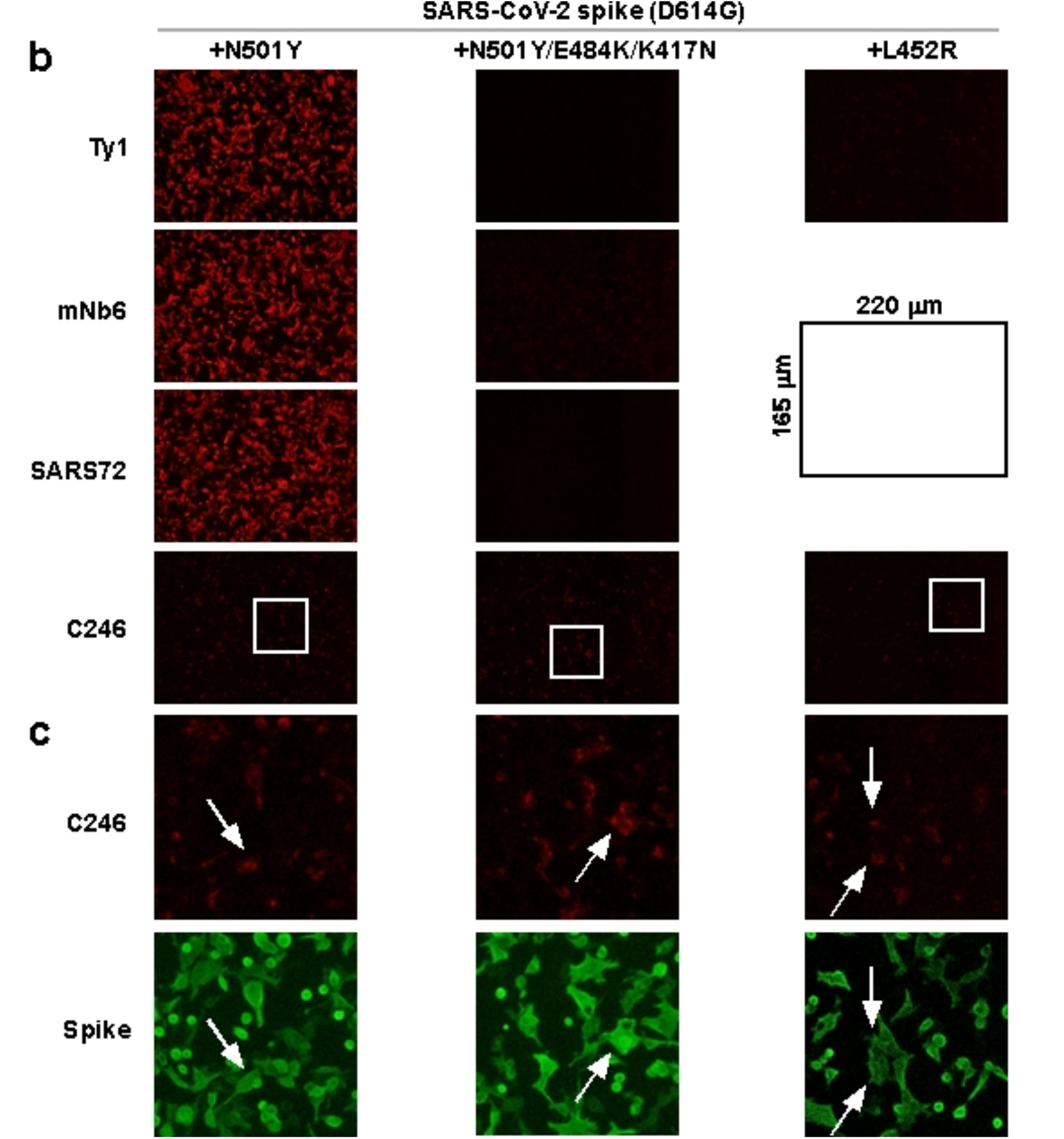
Microscopy analyses. **a,b**, Photographs of stained HEK cells expressing the SARS-CoV-2 spike variants carrying mutations in the RBD, original (D614G); alpha (N501Y); beta (K417N, E484K, and N501Y); and delta (L452R), with generated (**a**) and previously reported (**b**) nanobodies. **c**, Zoomed-in views of regions highlighted with white squares in (**b**): C246 intracellularly stained all spike variants without background—indicated by white arrows.

**Extended Data Fig. 5.**
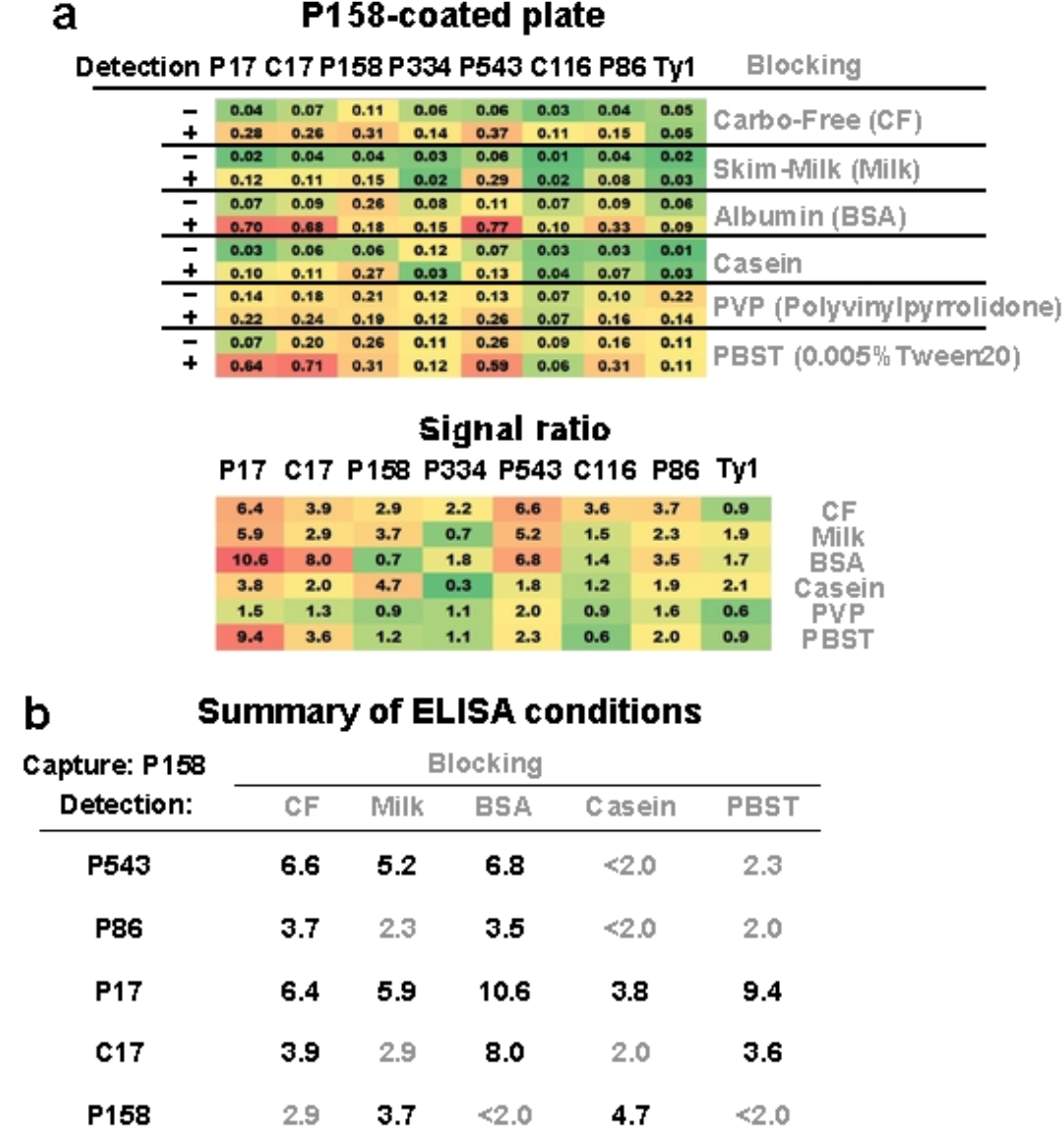
ELISA condition screenings. **a**, P158 was coated on an ELISA plate as a capture antibody; five kinds of blocking solutions (grey)—Carbo-Free (CF), skim milk (Milk), bovine serum albumin (BSA), casein, polyvinylpyrrolidone (PVP), and nonblocking saline (PBST)—were tested. Signals from 2 μg of the purified SARS-CoV-2 spike (+) or blocking buffer only (–) were compared among eight clones used as detection antibodies. Measured optical densities (OD=450 nm) and signal ratios (relative changes in OD_450 nm_ with (+) compared to without the spike protein (–)) are gradually coloured from green (low) to red (high). **b**, Summary of the top five detection antibodies and appropriate blocking conditions for the P158-based sandwich ELISA: ratios >3.0 are shown in black.

**Extended Data Fig. 6.**
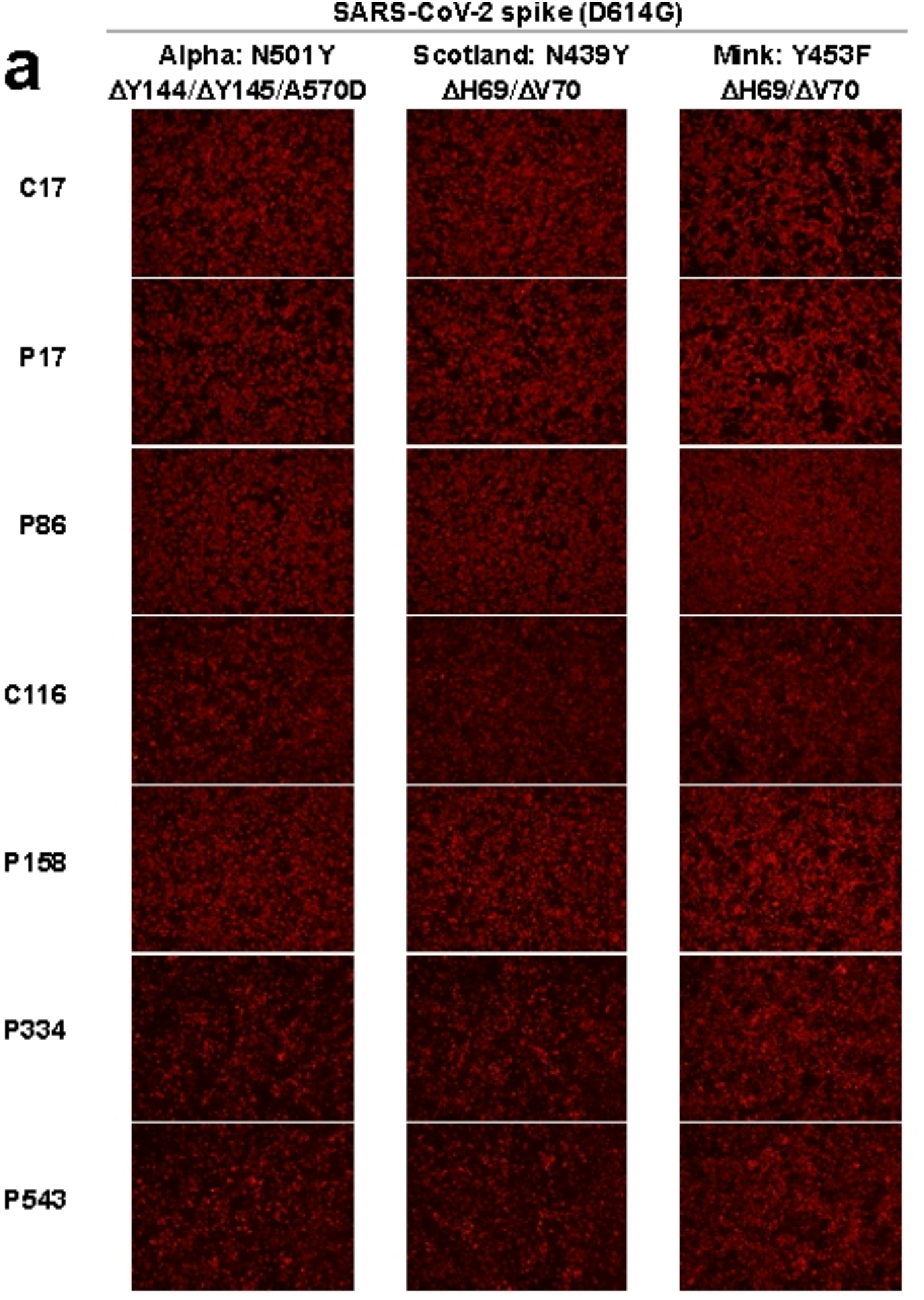
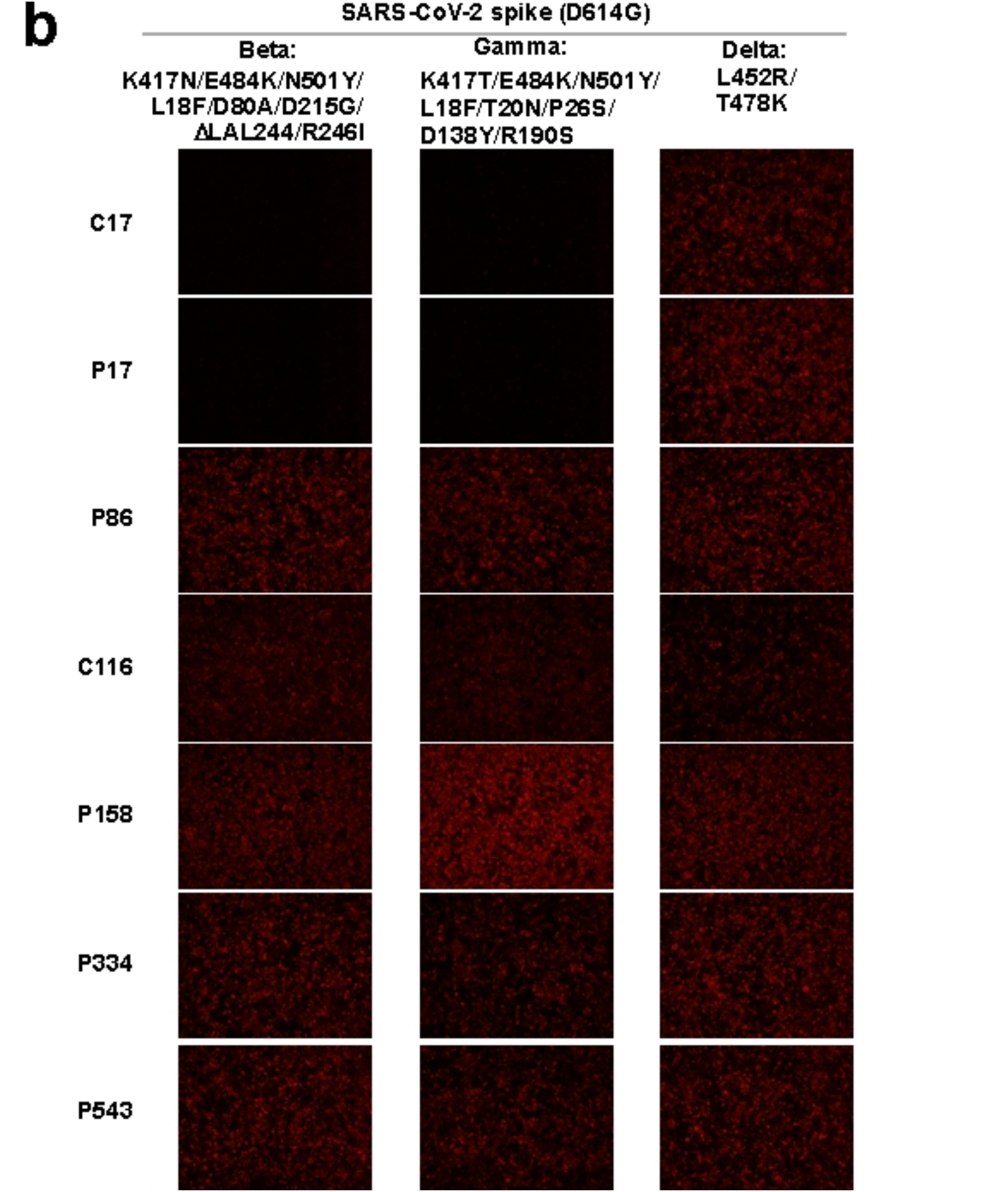
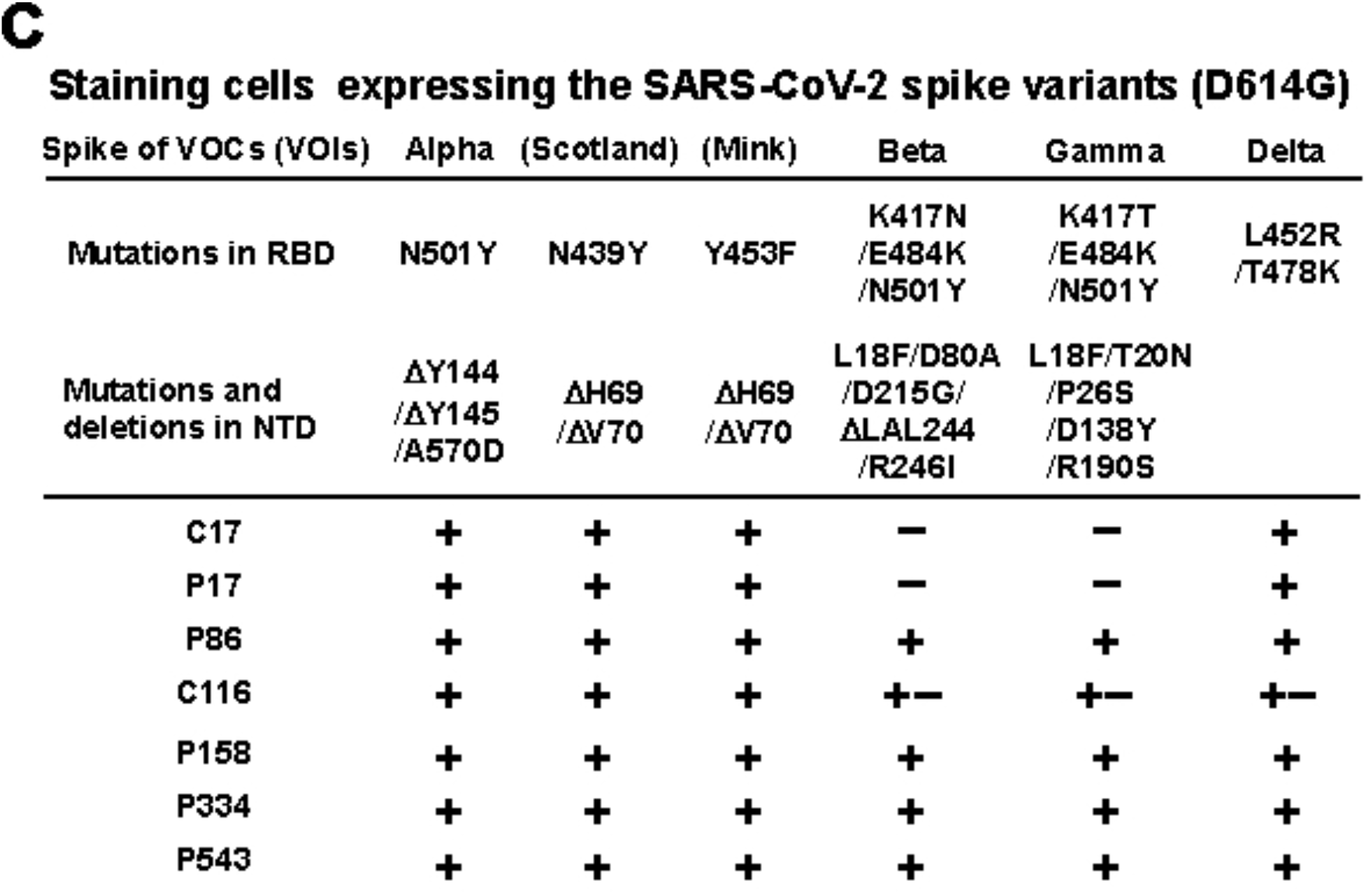
Microscopy analyses. Photographs of stained HEK cells expressing the SARS-CoV-2 spike (D614G) variants carrying mutations in the S1 domain: **a,b**, alpha (N501Y, Y144del, Y145del, and A570D); Scotland (N439Y, V69del, and H70del); Danish mink (Y453F, V69del, and H70del); beta (K417N, E484K, N501Y, L18F, D80A, D215G, L242del, A243del, L244del, and R246I); gamma (K417T, E484K, N501Y, L18F, T20N, P26S, D138Y, and R190S); and delta (L452R and T478K). **c**, Results of microscopy analyses are summarized: +) positive; +–) weak; or –) negative. The SARS-CoV-2 spike variants of interest (VOIs)—Scotland and Danish mink—are shown in parentheses.

**Extended Data Fig. 7.**
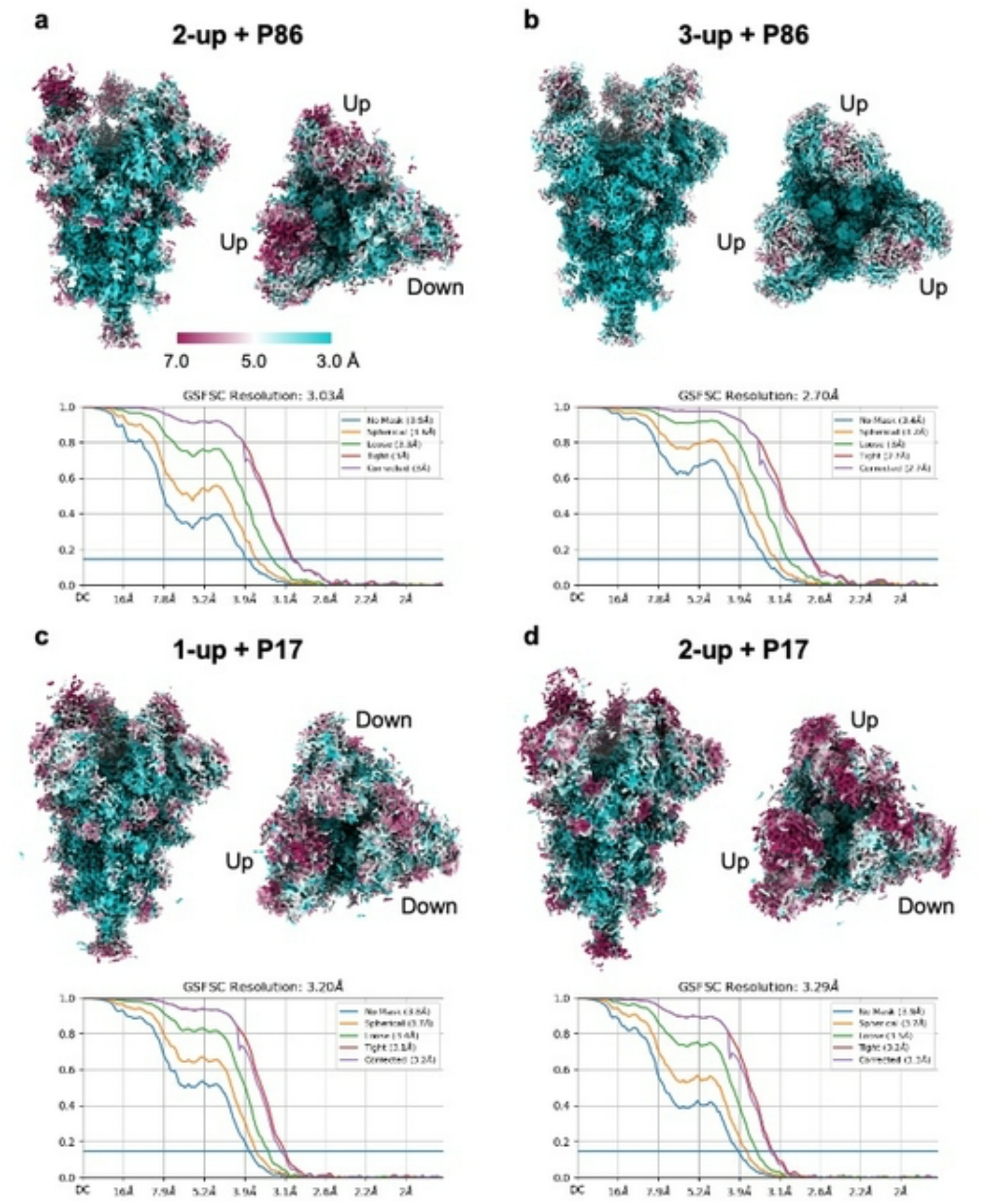
Cryo-EM density maps of spike-nanobody complexes. **a-d**, Upper panels show final sharpened maps of 2-up+P86 (**a**), 3-up+P86 (**b**), 1-up+P17 (**c**), and 2-up+P17 (**d**) datasets from side views (left) and top views (right) coloured by local resolutions (3.0–7.0 Å). Lower panels show the Fourier shell correlation (FSC) curves for the corresponding final maps of the datasets.

**Extended Data Fig. 8.**
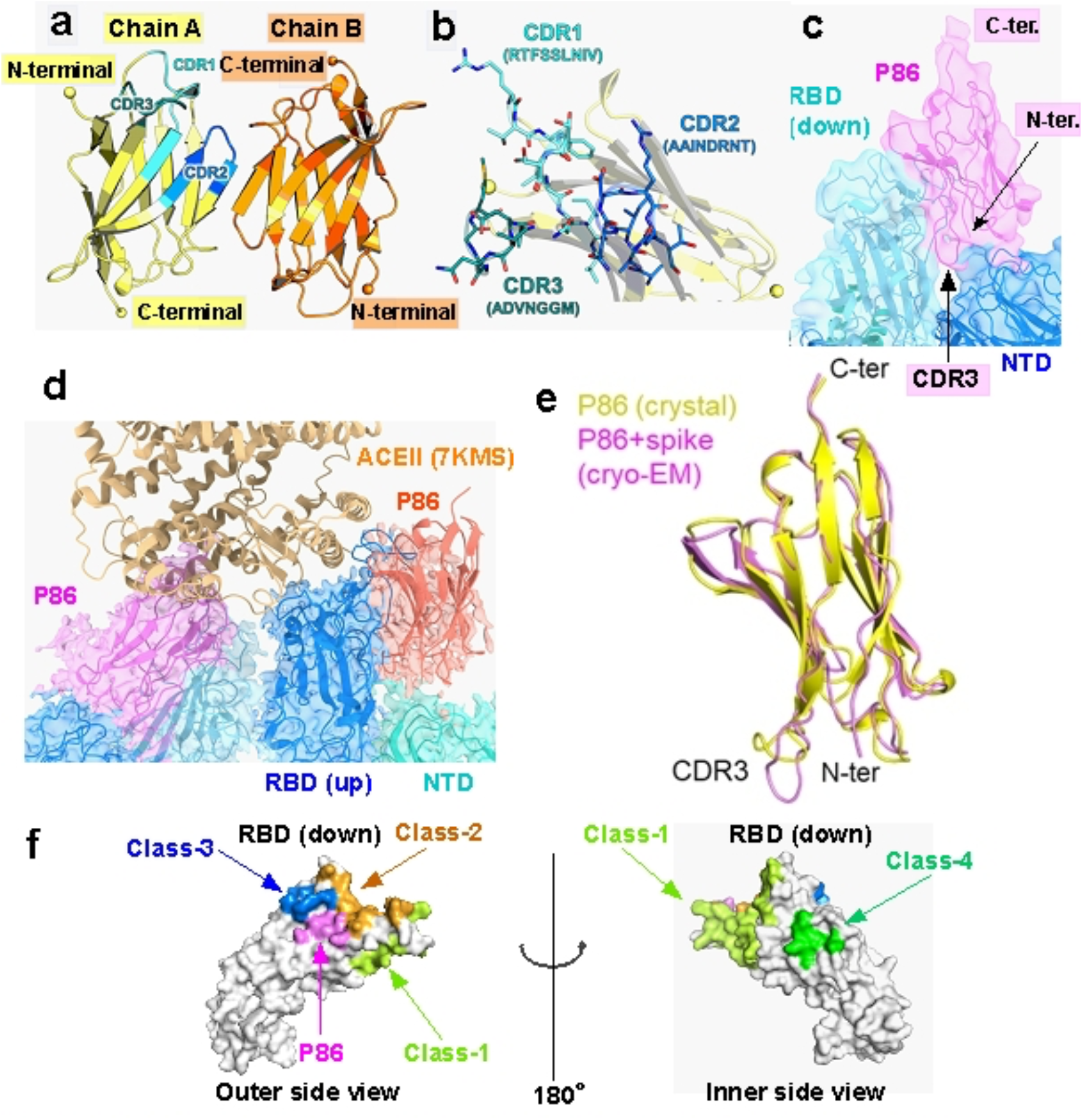
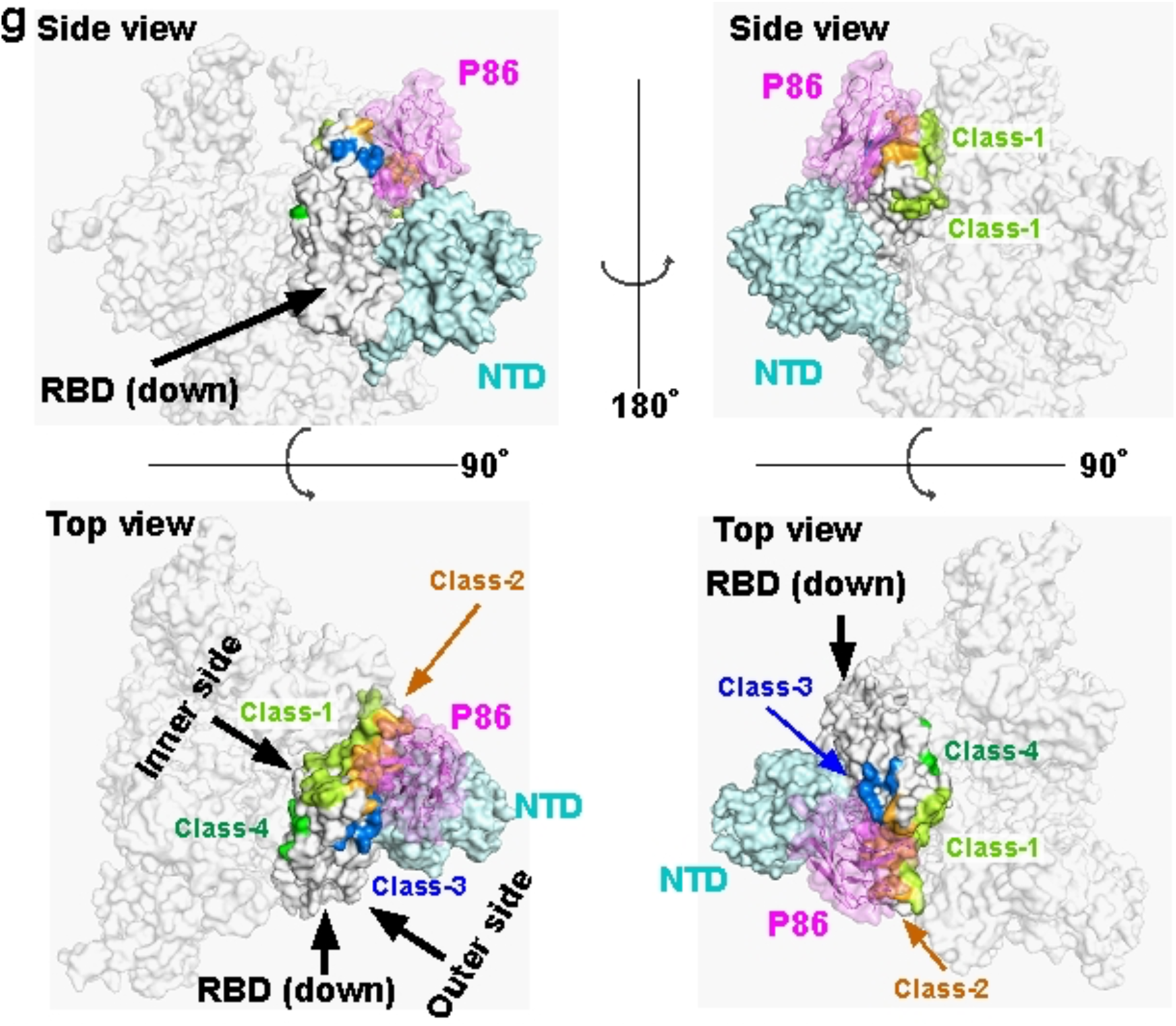
Crystal structure of P86 and a binding model of spike-P86 complex. **a**, Two P86 monomers in the asymmetric unit. Chain A and Chain B are almost identical, with a root mean square deviation for C_α_ atoms of 0.20 Å. **b**, Close-up view of three CDR regions in Chain A. **c**, Sharpened map from local refinement (5.13 Å resolution at FSC=0.143) around P86 bound to the down-RBD with our fitted model. C-terminal (C-ter.) and N-terminal (N-ter.) ends are indicated. **d**, Close-up view around P86 bound to the up-RBD with our fitted model structure. The model of ACE2 bound to the spike trimer (PDB entry: 7KMS) is superposed on the up-RBD shown in blue. **e**, Structure comparison between P86 alone (crystal structure) and P86 complexed with spike trimer (cryo-EM). **f**, Epitope mapping on the down-RBD. The epitopes of Class-1 to Class-4 compared to P86. **g**, A fitting model of P86 and the SARS-CoV-2 spike trimer. The down-RBD, the neighbouring NTD, and P86 are coloured white, cyan, and pink, respectively; other domains and protomers of the SARS-CoV-2 spike trimer are in reduced transparency.

**Extended Data Table 1.** Cryo-EM data collection and processing.

| Dataset | 2-up+P86<br>(EMDB-32078)<br>(PDB ID: 7VQ0) | 3-up+P86<br>(EMDB-32079) | 1-up+P17<br>(EMDB-32080) | 2-up+P17<br>(EMDB-32081) |
| --- | --- | --- | --- | --- |
| **Data collection and processing** |  |  |  |  |
| Magnification | 60,000 | 60,000 | 60,000 | 60,000 |
| Voltage (kV) | 300 | 300 | 300 | 300 |
| Electron exposure (e<sup>-</sup>/Å<sup>2</sup>) | 60 | 60 | 60 | 60 |
| Defocus range (μm) | -0.5 to -2.0 | -0.5 to -2.0 | -0.5 to -2.0 | -0.5 to -2.0 |
| Pixel size (Å) | 0.870 | 0.870 | 0.874 | 0.874 |
| Symmetry imposed | C1 | C3 | C1 | C1 |
| Imported movies (no.) | 4,175 | 4,175 | 2,124 | 2,124 |
| Initial particle images (no.) | 878,975 | 878,975 | 766,652 | 766,652 |
| Final particle images (no.) | 38,004 | 44,292 | 67,529 | 48,715 |
| Map resolution (Å) | 3.03 | 2.70 | 3.20 | 3.29 |
| FSC threshold | 0.143 | 0.143 | 0.143 | 0.143 |
| **Refinement** |  |  |  |  |
| Model vs. Map CC (volume) | 0.79 |  |  |  |
| Protein residues | 3624 |  |  |  |
| R.m.s. deviations |  |  |  |  |
| Bond lengths (Å) | 0.003 |  |  |  |
| Bond angles (°) | 0.516 |  |  |  |
| **Validation** |  |  |  |  |
| MolProbity score | 1.81 |  |  |  |
| Clashscore | 9.09 |  |  |  |
| Rotamer outliers (%) | 0.03 |  |  |  |
| **Ramachandran plot** |  |  |  |  |
| Favored (%) | 95.30 |  |  |  |
| Allowed (%) | 4.61 |  |  |  |
| Disallowed (%) | 0.08 |  |  |  |

**Extended Data Table 2.** Crystallographic data collection and refinement statistics.

| **Data Collection** |  |
| --- | --- |
| X-ray source | SPring-8 BL44XU |
| Wavelength (Å) | 0.9000 |
| Space group | *C2* |
| Unit cell *a*, *b*, *c* (Å) | 105.08, 43.17, 56.03 |
| Unit cell *α*, *β*, *γ* (°) | 90.00, 110.76, 90.00 |
| Resolution range (Å) | 49.13–1.60 (1.63–1.60)* |
| Total No. of reflections / No. of unique reflections | 103,494 (5,156) / 30,851 (1,493) |
| Completeness (%) | 99.1 (99.5) |
| Redundancy | 3.4 (3.5) |
| $\langle I/\sigma(I) \rangle$ | 11.9 (2.8) |
| *R*<sub>meas</sub> (all I+ & I-) | 0.067 (0.668) |
| *R*<sub>meas</sub> (within I+/I-) | 0.065 (0.628) |
| CC<sub>1/2</sub> | 0.998 (0.835) |
| **Refinement** |  |
| Resolution range (Å) | 30.80–1.60 (1.65–1.60) |
| Completeness (%) | 98.8 (99.0) |
| No. of reflections, working set | 30,833 (3,054) |
| No. of reflections, test set | 1,510 (138) |
| *R*<sub>work</sub> / *R*<sub>free</sub> | 0.177/0.213 (0.236/0.245) |
| No. of non-H atoms | 2167 |
| Protein | 1885 |
| Ion/Ligand | 54 |
| Water | 232 |
| R.m.s. deviation bonds (Å), angles (°) | 0.006, 0.87 |
| Average *B* factors (Å<sup>2</sup>) | 23.61 |
| Protein | 22.29 |
| Ion/Ligand | 34.89 |
| Water | 31.84 |
| Ramachandran favored / allowed / disallowed (%) | 98.25 / 1.75 / 0 |
| PDB code ID | 7VPY |

\*Values in parentheses refer to the highest resolution shell.

